# p63 regulates stem cell maintenance and age-associated functional decline in human airway basal cells

**DOI:** 10.64898/2026.07.22.740085

**Authors:** Andrew S. Farr, Jessica C. Orr, Buthainah M. Ahmed, Léa Hascher, Tony Brooks, Robert E. Hynds

**Author notes:** **Correspondence:** Robert E. Hynds.

## Abstract

Age is a principal risk factor for chronic respiratory diseases. During aging, the airway epithelium undergoes structural and functional changes, including a reduced regenerative capacity. Basal cells act as stem/progenitor cells within the airway epithelium and are known to acquire age-dependent intrinsic defects, such as genomic mutations, transcriptomic changes and declining stem cell potential. However, the molecular mechanisms bridging transcriptomic change and stem cell function are unknown. Here we show that the transcription factor p63, a key regulator of epithelial cell identity, controls airway basal cell progenitor function *in vitro* and declines in expression with aging. We found impaired *in vitro* 2D colony and 3D organoid formation potential, as well as reduced culture longevity in primary human nasal basal cells isolated from older adults (>70 years) compared to those from pediatric (<10 years) donors. p63 protein expression was lower in basal cells from older adults and p63-regulated genes were enriched among genes whose expression changed with aging. Overexpression of p63 in older adult basal cells partially rescued progenitor cell functions, while gene knockdown in pediatric cells caused functional deficits. Integrative transcriptomic analyses revealed the network of p63-regulated genes in basal cells and demonstrated similarities in pathway engagement between *TP63*-knockdown cells and aged epithelium. Thus, while exogenous p63 expression alone could not rejuvenate older adult epithelial cells, p63 has an influential role in declining epithelial stem cell function during airway aging. Defining the upstream mechanisms controlling p63 decline during epithelial aging may reveal novel targets to promote healthy aging and reduce the burden of chronic lung disease.

## Introduction

Aging is a primary risk factor for chronic respiratory diseases, which are leading causes of morbidity and mortality worldwide^1^. Accumulating evidence indicates that the core cellular hallmarks of aging converge on pathways involved in tissue maintenance, regeneration and repair^2^, implicating age-related dysfunction of these processes in disease pathogenesis. In the airway epithelium, these functions are driven by basal stem/progenitor cells and age-dependent changes in these cells likely drive epithelial dysfunction and disease susceptibility.

Several lines of evidence point to mechanisms through which airway epithelial cell aging may contribute to respiratory disease risk. The accumulation of somatic mutations in airway basal cells with age contributes to the early evolution of lung squamous cell carcinoma^3,4^, implicating intrinsic alterations in basal cells in disease pathogenesis. Moreover, age-associated accumulation of senescent cells and the accompanying senescence-associated secretory phenotype (SASP) can have pro-inflammatory and pro-fibrotic effects in the airway^5^, demonstrating that age-related epithelial changes drive wider microenvironmental dysregulation. Functionally, air-liquid interface (ALI) cell cultures derived from basal cells of older adults exhibit altered responses to viral infection, with more viral replication following SARS-CoV-2 infection^6^ and a dysregulated immune response following influenza infection^7^. These observations collectively point to basal cell dysfunction as a feature of airway epithelial aging.

As long-lived tissue-resident stem/progenitor cells, airway basal cells are positioned to integrate chronic environmental and inflammatory exposures (such as tobacco smoke, air pollution, and infections), leading to the cumulative accrual of detrimental genetic and epigenetic alterations and consequent functional defects over the life course. Pediatric epithelial cells have distinct transcriptional^8^ and proteomic^9^ profiles compared to those from adults, consistent with altered stem cell capacity and supported by their greater longevity in cell culture^8^. However, beyond these observations, the key mechanisms remain undefined, and understanding of how airway basal cell phenotype and function change with advanced age is limited.

The transcription factor tumor protein p63 (encoded by *TP63*) is highly expressed in basal cells across stratified and pseudostratified epithelia, including the airway^10–15^. *TP63* is expressed from two transcription start sites, namely the TA and ΔN promoters, and is subject to alternative splicing that gives rise to at least five splice variants (α, β, γ, δ, ε). Although studies of p63 function in airway epithelial cells specifically are few in number, *TP63* knockdown in immortalized human bronchial cell lines^16,17^ and primary mouse tracheal cells^17,18^ are consistent with findings in other tissues. These studies indicate that ΔNp63 isoforms regulate multiple aspects of the basal stem cell phenotype^19^, including proliferation^15,20–23^, adhesion^14^, self-renewal and differentiation^21,22,24–26^.

p63 has been implicated in regulating multiple developmental signaling pathways (including Notch, Wnt/β-catenin, Hedgehog, and Hippo pathways^27–31^), as well as chromatin architecture^32^. Tissue context is likely to be important: while some canonical p63 targets, such as the NOTCH ligand JAG1, are common across epithelia, profiling in skin cells^33–37^, prostate cells^38^ and cancer cell lines^24,39–42^ has demonstrated both tissue- and context-specific differences in p63 binding and regulatory activity. In cultured primary human airway basal cells, chromatin immunoprecipitation sequencing (ChIP-seq) identified overlap in p63, YAP and TEAD genomic binding sites that suggested a regulatory complex associated with driving airway basal cell proliferation and immune cell communication^28^. Beyond this, the specific functions of p63, its gene regulatory network and factors that regulate these in airway basal cells remain poorly understood.

In this study, we set out to define intrinsic changes to airway epithelial stem cells during aging using a primary cell culture approach. We assessed *ex vivo* stem cell phenotype and function in nasal epithelial basal cells from children and older adults, identifying reduced *TP63* expression in older adult cell cultures undergoing age-related functional decline. Gene knockdown in pediatric basal cells impacted stem cell functions, while overexpression partially recovered function in older adult basal cells. Together, these data identify p63 as a regulator of airway epithelial stem cell aging.

## Results

To investigate age-dependent functional differences in airway basal cells, we collected nasal brush biopsies from participants who were either under 10 years of age (pediatric) or over 70 years of age (older adults). We cultured these in co-culture with mitotically-inactivated 3T3-J2 mouse embryonic fibroblasts in medium containing the small molecule inhibitors Y-27632 and WS6 to enable expansion of nasal epithelial cells from the few basal cells present in initial biopsy samples (**Fig. 1A**; ^43^). We generated passagable cell cultures from 28 of 29 pediatric donors and 8 of 12 older adult donors (**Fig. 1B**). Four donor cell cultures from each age group were characterized over 16 passages. Over long-term culture, pediatric basal cells had a greater capacity for expansion, with all four older adult cultures and one pediatric culture terminating at or before passage 16 (**Fig. 1C**). After brief *in vitro* expansion, the colony formation potential of pediatric basal cells was significantly higher than older adult basal cells, suggesting that a greater proportion of basal cells from pediatric donors had colony forming potential (**Fig. 1D**). Pediatric basal cells maintained their colony forming ability over 16 passages, albeit at a reduced level compared to passage 4 cultures, while older adult basal cells consistently lost clonogenicity over time in culture (**Fig. 1E**). Organoid forming ability followed a similar trend to the colony formation assays, with preserved organoid formation in pediatric but not older adult donors at passage 16 (**Fig. 1F**), although both pediatric and older adult basal cells were able to form organoids of comparable size (**Supplementary Fig. 1**) containing differentiated mucosecretory and ciliated cells, including at late passage (**Fig. 1G**).

**Figure 1:**
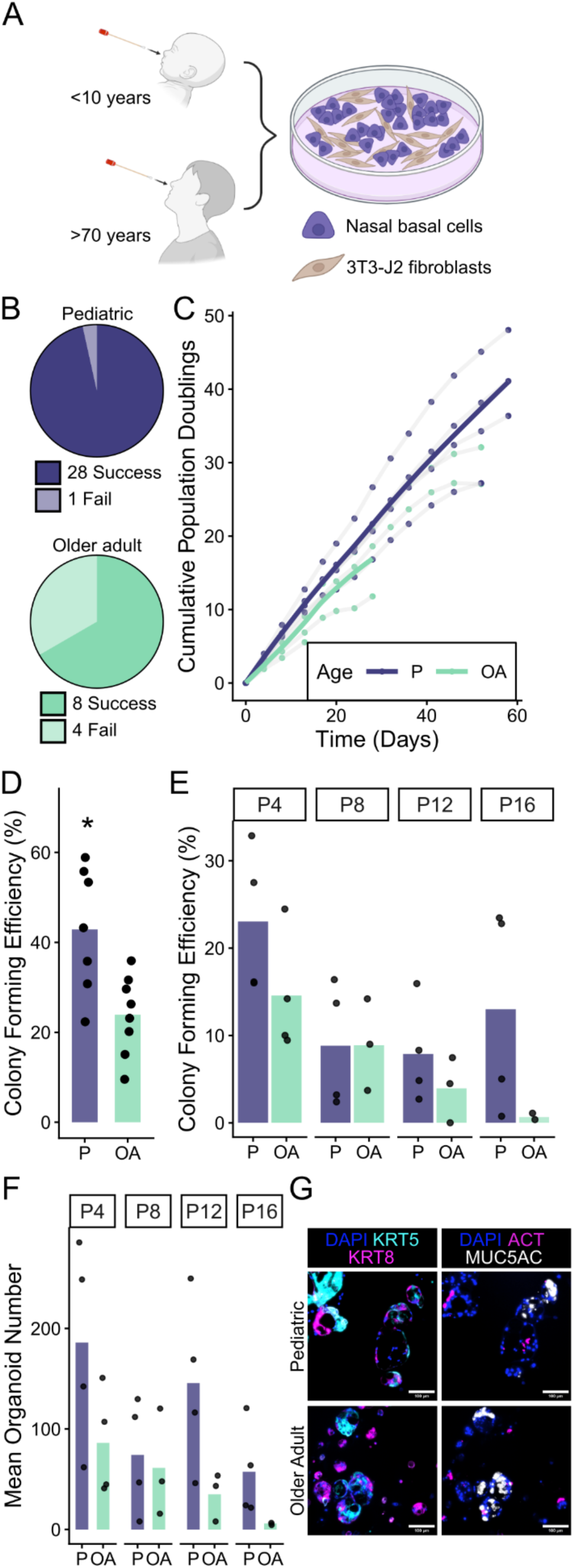
Age-associated functional decline in primary human nasal epithelial basal cells. A) Nasal basal cells from pediatric (P) and older adult (OA) donors were obtained by nasal brush biopsy and expanded in 3T3-J2 fibroblast co-culture. B) Proportion of nasal brushings from P (upper) and OA (lower) donors that produced passagable basal cell cultures. C) Population doublings for P and OA basal cells from one passage after thawing (passage four post explant; n = 4 donors per age group). D) Bars show mean colony forming efficiency (CFE) for each age group, with points indicating the mean of three technical well replicates per donor (unpaired t-test; n = 7 P, 8 OA donors). E) CFE at passages 4, 8, 12 and 16. Bars show mean CFE for each age group with points indicating the mean of three well replicates per donor (Wilcoxon signed-rank tests; n = 4 P, 2-4 OA donors; absence of stars indicates non-significance). F) Organoid number from P or OA basal cells at passage 4, 8, 12 or 16 (Wilcoxon signed-rank test; n = 4 P, 2-4 OA donors, points show the mean of 8 technical well replicates per donor; lack of stars indicates non-significance). G) Representative immunofluorescence staining of organoids derived from passage 12 P or OA basal cells. Staining shows basal-luminal polarization (left: KRT8, magenta; KRT5, cyan; DAPI, blue) and presence of mucosecretory and multiciliated cells (right: MUC5AC, gray; ACT, magenta; DAPI, blue). Scale bars = 100 µm.

Bulk transcriptomic analysis of six pediatric and six older adult donor cultures at early passage identified 878 genes significantly differentially expressed by at least a 1.5-fold change (**Fig. 2A**). A greater number of these genes were overexpressed in the older adult donor cells (561 in older adult compared with 307 in pediatric; **Fig. 2A**). Gene set enrichment analysis (GSEA) with MSigDB hallmark gene sets identified 17 significantly enriched pathways within the dataset (**Supplementary Fig. 2**). Consistent with enhanced proliferative capacity, G2M checkpoint, E2F and MYC pathways were enriched in pediatric cells (**Fig. 2B**). In contrast, key pathways enriched in older adult cells were associated with responses to stress and immunological activity (**Fig. 2B**).

**Figure 2:**
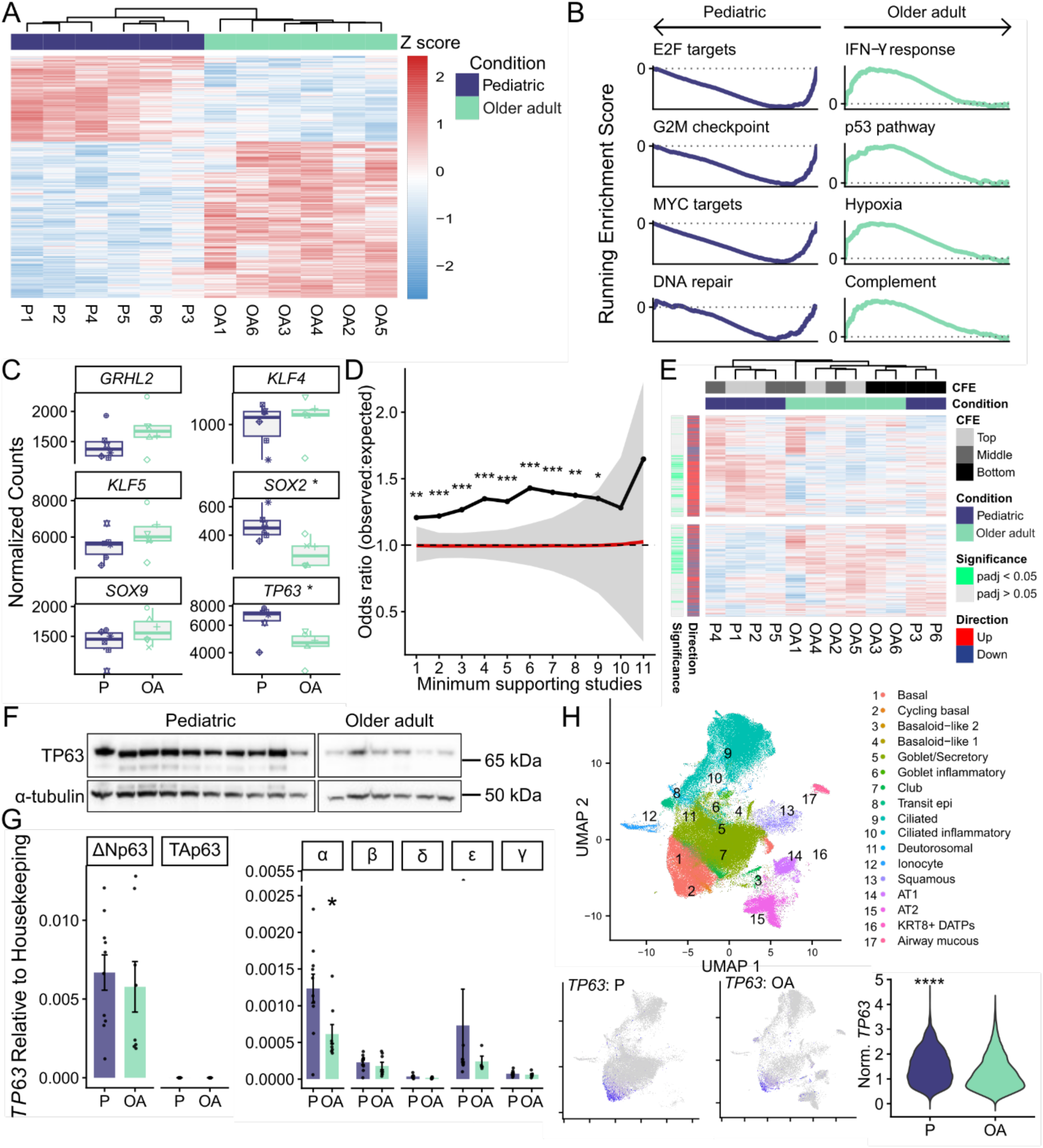
Age-associated transcriptomic differences implicate p63 as a regulator of basal cell functional decline. **A)** Differentially expressed genes (log2 fold change > 0.58 and adjusted p-value < 0.05) between pediatric (P) and older adult (OA) donor cell cultures at passages 3-4 in bulk RNA sequencing. Variance-stabilized expression values were used for visualization. Hierarchical clustering of samples was performed using Euclidean distance. **B)** Selected pathways significantly enriched (p < 0.05) within P (left) and OA (right) in transcriptomes, identified by Gene set enrichment analysis. Curves show running enrichment scores. **C)** Normalized gene expression values for selected transcription factors, identified in the facet titles. Boxplots show median and interquartile range of expression in P and OA groups, points represent donors. Statistical significance was determined by differential expression analysis in DESeq2 with a Wald test and Benjamini-Hochberg multiple testing correction (n = 6 donors per age group). **D)** Enrichment of p63-regulated genes among differentially expressed genes between age groups using a Fisher’s exact test. A positive odds ratio indicates enrichment. Odds ratios are plotted for genes with p63-dependent expression supported by 1-11 previous studies. Gray cloud shows the 95% confidence interval from 5,000 random permutations and the red line shows the mean of these permutations. Empirical p-value reflects the proportion of permutations exceeding the observed enrichment (n = 6 donors per age group). **E)** Heatmap showing expression of genes with p63-dependent expression supported by at least three studies. Unsupervised hierarchical clustering by Euclidean distance of samples was performed. Differentially expressed genes with an adjusted p-value < 0.05 and the direction of regulation by p63 are indicated by colored bars (left). Passage three colony formation efficiency (ranked as the top, middle, and bottom thirds within each age group) and the age group are also indicated by colored bars (top). **F)** Western blot using a pan-p63 antibody in passage two basal cells, with an α-tubulin loading control (n = 10 P, 6 OA donors). **G)** RT-qPCR quantification of *TP63* isoforms in passage two basal cells. Primers specific to all splice isoforms of the TA or ΔN transcription start sites (left), or specific isoforms (α, β, δ, ϒ or ε) from either transcription start site (right). RNA expression was plotted relative to the mean of *GAPDH* and *RPS13* housekeeping genes (unpaired t-tests; n = 10 P, 8 OA donors). **H)** Precomputed UMAP of a single cell RNA sequencing dataset from the airway epithelium of healthy P and OA donors (upper). Cell type annotations provided in the metadata were used to label clusters. *TP63* expression in P (lower left) and OA (lower middle) donors was visualized using feature plots with quantile-based cutoffs (q10–q90). *TP63* expression was compared in basal cells with detectable *TP63* expression (>0; Wilcoxon rank-sum test, lower right; P: n = 6,506 cells, OA: 4,591 cells).

We next investigated whether transcription factors known to regulate airway epithelial development (grainyhead-like transcription factor 2, *GRHL2*^44^, Kruppel-like factor 4, *KLF4*^45^), *KLF5*^45^, SRY-box transcription factor 2, *SOX2*^46^, *SOX9*^47^ and *TP63*^48^) were relevant to the altered gene expression observed in aging. Expression of *SOX2* and *TP63* were significantly downregulated in older adult donor cells (1.40-fold change in *SOX2* and 1.35-fold change in *TP63*; **Fig. 2C**). Initial ChIP-X enrichment analyses^49^, which are limited by the variable tissues, p63 isoforms and disease states investigated in the underlying studies, suggested potential enrichment of *TP63*, but not *SOX2*, targets within the age-associated differentially expressed genes (**Supplementary Fig. 3**). To investigate enrichment with a higher confidence list of p63 target genes, a recent meta-analysis of p63-regulated gene expression from 16 p63 expression modulation datasets^50^ was used. Fisher’s exact tests showed that genes with literature support for displaying p63-regulated expression were significantly enriched within the age-associated differentially expressed genes (**Fig. 2D**). Enrichment increased with increasing stringency in the cut-off for number of supporting studies between one and nine supporting studies, after which the number of target genes became small relative to the background (≤ 72 of 11,058 genes; **Fig. 2D**). In unsupervised clustering of the expression of literature-supported p63-regulated genes, the samples clustered by function, rather than by age. The four pediatric donor samples with the higher colony forming efficiency formed a separate cluster from the six older adult cultures and the two pediatric cultures with the lowest colony forming efficiency (**Fig. 2E**).

These transcriptional analyses suggested that age-associated changes to the expression of *TP63* and subsequent changes to the downstream p63 gene regulatory network could contribute to the functional deficiencies observed in the older adult donor cell cultures. Decreased p63 expression in older adult basal cells was confirmed at the protein level by Western blot analysis of early passage cell cultures (**Fig. 2F**). Isoform-specific qPCR demonstrated that the ΔNp63α splice variant is the dominant isoform in airway basal cells and also showed the greatest age-associated difference in RNA expression (**Fig. 2G**). To validate this finding *in vivo*, the expression of *TP63* and *SOX2* were examined in a single cell RNA sequencing dataset of airway epithelial cells from fresh brushings of healthy pediatric and older adult donors^6^. In total, 191,600 cells were contained in the dataset (112,708 pediatric, 78,892 older adult) from 93 donors (49 pediatric, 44 older adult)^6^. Visualization on UMAP plots showed that *TP63* expression was concentrated in the basal and cycling basal cell types (**Fig. 2H**), while *SOX2* expression was found across airway epithelial cell types (**Supplementary Fig. 4**). Among basal cells expressing the respective gene, *TP63* and *SOX2* showed significantly greater expression levels in pediatric compared to older adult cells (1.34-fold change in *TP63* and 1.28-fold change in *SOX2*), consistent with *in vitro* data (**Fig. 2H; Supplementary Fig. 4**).

To investigate whether restoring p63 expression could drive pediatric-like basal cell function in older adult cells, we used a doxycycline-inducible lentiviral approach (**Fig. 3A**). Cells were transduced with two lentiviruses; one containing a vector with the ΔNp63α coding sequence (NM_001114980) under the control of a tetracycline response element (TRE) doxycycline-inducible promoter (**Fig. 3A**; *ΔNp63α*-mCherry vector), and another containing the reverse tetracycline transactivator (rtTA) protein under the control of a CMV promoter (**Fig. 3A**; Tet-On vector). The dual-vector system facilitated doxycycline-inducible overexpression of *ΔNp63α*, in a concentration-dependent manner (**Supplementary Fig. 5**). However, a high doxycycline concentration significantly reduced colony formation in untransduced cells (**Supplementary Fig. 5**). Six older adult cell cultures were transduced with both vectors in two experimental replicates. Five out of six donor cultures could survive the selection process after transduction (10 out of 12 independent transductions) and in all five donors the addition of doxycycline (0.5 µg/mL) led to robust overexpression of *ΔNp63α* at the protein level (**Fig. 3B**). The low baseline expression of p63 (**Fig. 3B**) was consistent with low endogenous p63 expression previously observed in older adult cells (**Fig. 2F**) and Western blot with a pan-p63 antibody confirmed that only the *ΔNp63α* isoform was being overexpressed (**Fig. 3B**).

**Figure 3:**
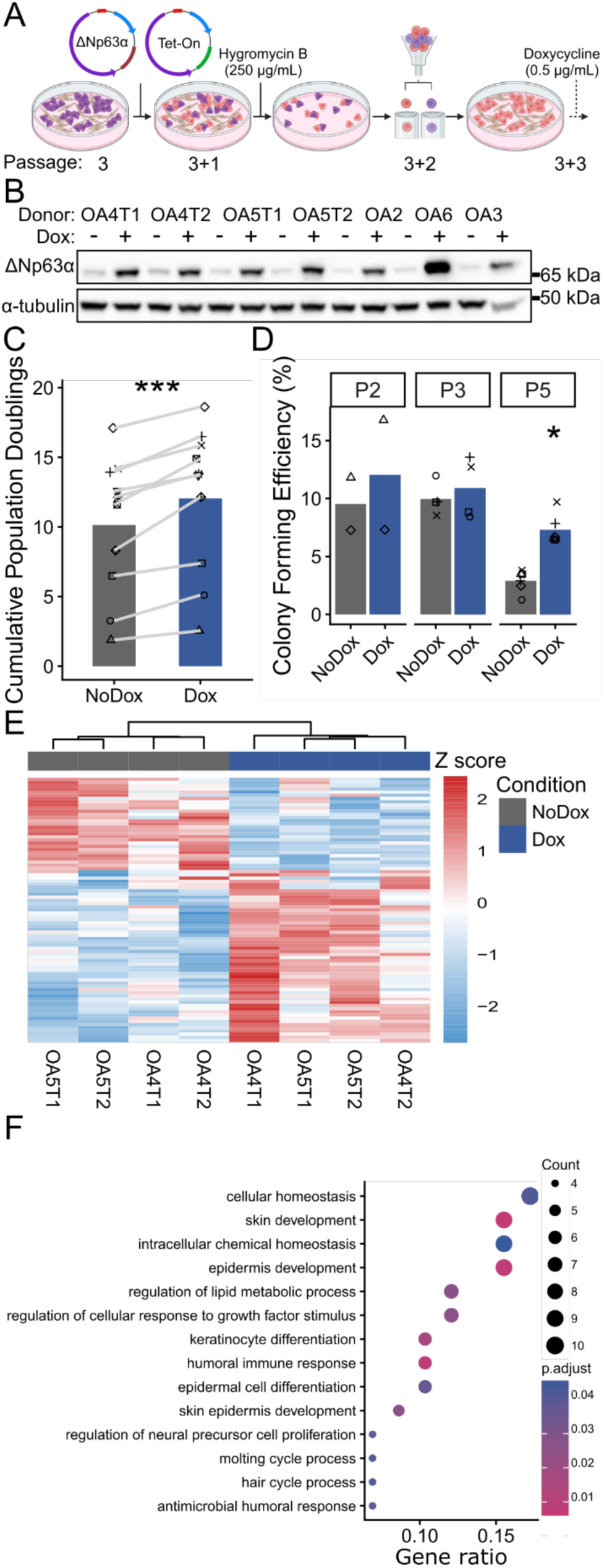
ΔNp63 overexpression improves aspects of stem cell function in primary older adult basal cells. **A)** Schematic representation of the overexpression experimental design. **B)** Western blot showing ΔNp63α protein expression in primary older adult (OA) donors transduced with the *TP63* overexpression vectors after one passage with or without doxycycline induction (n = 7 independent transductions of 5 donor cell cultures). Protein lysates were collected three passages after transduction (PT+3). α-tubulin was used as a loading control. **C)** Cumulative population doublings for OA basal cells transduced with both *TP63* overexpression vectors, divided into +/- doxycycline conditions and cultured until replicative senescence. A paired t-test was used to compare the mean final cumulative population doublings reached by each donor between conditions (n = 10 independently transduced cell cultures, 5 donors). **D)** Colony formation efficiency of primary OA basal cells transduced with *TP63* overexpression vectors and divided into +/- doxycycline conditions. Assays were plated after two, three or five passages (indicated by facet labels). For Facets 2 and 3, paired Wilcoxon signed-rank tests were performed (n = 4-6 independent transductions of 2 donors). No statistical test was performed for Facet 1. **E)** Differentially expressed genes (log2 Fold Change > 0.58 and adjusted p-value < 0.05) between transduced OA cultures divided into +/- doxycycline conditions for five passages (condition indicated by colored bar, top). Hierarchical clustering of samples was performed using Euclidean distance. **F)** Gene Ontology (GO) analysis of genes with a log2 fold change > 0 (trend upregulated after *TP63* overexpression).

Two transduced cell cultures (OA4 and OA5) were capable of maintaining long-term proliferation after the selection process and therefore three independent transductions of each donor were functionally characterized over five passages after selection. Population doublings within this time were comparable between conditions until day 20, at which point the proliferation of control cells slowed and ceased by Day 30 (**Fig. 3C**). Overexpression of *ΔNp63α* led to a modest but statistically significant increase in population doubling potential, with proliferation ceasing one or two passages later than controls in all cell cultures (**Fig. 3C**). Colony formation assays across multiple passages identified a significant improvement in *ΔNp63α*-overexpressing cells (**Fig. 3D**). Between two and five passages after induction of overexpression (between seven and ten passages total) colony forming efficiency was close to 10% (**Fig. 3D**), comparable to the efficiency of pediatric donor cell cultures at equivalent total passage numbers (**Fig. 1C**). Differentiated air-liquid interface (ALI) cultures from either control or *ΔNp63α*-overexpressing cells demonstrated intact epithelial barrier function, as assessed by transepithelial electrical resistance (TEER; **Supplementary Fig. 6A**), but did not produce multiciliated cells (**Supplementary Fig. 6B**), presumably due to the overall high passage number of older adult cells, along with the inability of *ΔNp63α* overexpression to overcome the late passage loss of differentiation potential.

Consistent with partial rescue of basal cell progenitor functions, bulk RNA sequencing identified relatively few differentially expressed genes with over 1.5-fold change (53 up and 30 down; **Fig. 3E**). However, GO analysis of significantly differentially expressed genes identified pathways, including cellular homeostasis, epithelial development and response to growth factor stimulus, as being upregulated following *ΔNp63α* overexpression (**Fig. 3F**).

We next used constitutive lentiviral shRNA knockdown to assess the necessity of *TP63* expression for basal progenitor cell functions (**Fig. 4A**). Initially, four shRNA clones were tested for their effect on *ΔNp63* RNA expression and colony formation efficiency in one pediatric donor culture (**Supplementary Fig. 7A-B**). All four reduced both expression and colony formation efficiency (**Supplementary Fig. 7A-B**). Six pediatric donor cell cultures were transduced with either an empty vector or a vector containing shRNA targeting Exon 6 of the *TP63* DNA binding domain, which is conserved between all *TP63* isoforms (**Fig. 4A**). Transduction with shRNA resulted in knockdown of p63 at the protein level relative to empty vector-transduced controls in all six donor cell cultures (**Fig. 4B**). *TP63* knockdown significantly reduced the rate of long-term proliferation and capacity for population doublings (**Fig. 4C**). *TP63* knockdown caused a 24.6% mean reduction in cumulative population doublings over the course of 24 days in culture, with a high degree of variation between donors (EV = 12.4 (± 4.0) doublings, shRNA = 9.3 (± 3.0) doublings). In colony formation assays at passage 2+3 (cells expanded for three passages after transduction at passage two), knockdown reduced colony forming efficiency compared to empty vector in all donors (**Fig. 4D**). *TP63* knockdown resulted in a failure to proliferate past passage 2+7 in 3/6 donors (**Fig. 4C**), and colony formation assays on the remaining three cell cultures at passage 2+7 showed lower colony formation in *TP63* knockdown conditions (**Fig. 4D**). Although some colony forming ability was retained in the knockdown cells, fluorescence imaging showed that these colonies were formed by GFP-negative cells, suggesting that the colonies were formed by either contaminating untransduced cells or cells who had downregulated shRNA expression (**Supplementary Fig. 7C**).

**Figure 4:**
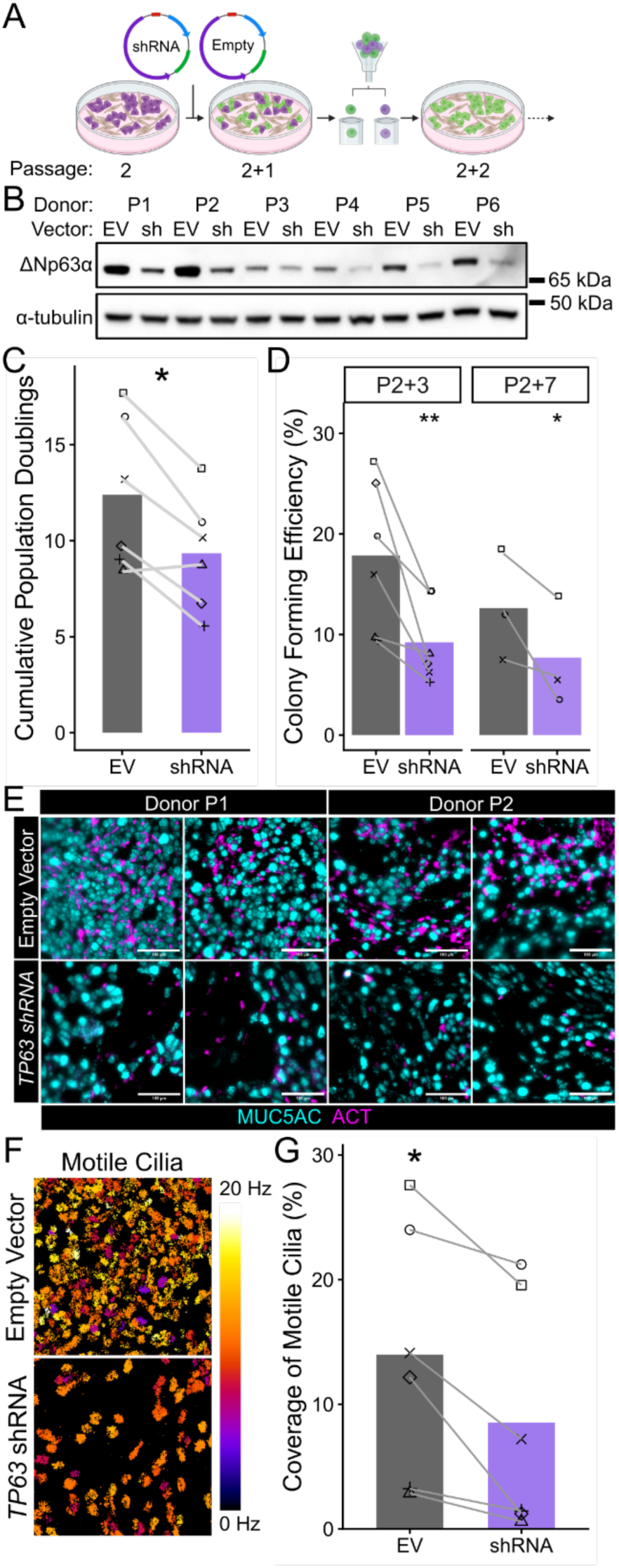
Knockdown of *TP63* in primary pediatric basal cells reduces progenitor cell function. Pediatric (P) donor cultures were transduced with either lentiviral shRNA against *TP63* or an empty vector (EV) construct. Transduced cultures were selected by fluorescence-activated cell sorting for GFP expression. Western blot of ΔNp63α protein expression in six primary pediatric donors transduced with either *TP63*-shRNA or EV control vectors. α-tubulin was used as a loading control. **C)** Cumulative population doublings for P basal cells transduced with either *TP63*-shRNA or EV control vectors (paired t-tests, n = 6 donors, indicated by shape). **D)** Colony forming efficiency three (n = 6 donors) or seven (n = 3 donors) passages after transduction. Paired Wilcoxon signed-rank tests were used to compare means of three technical well replicates per condition. **E)** Immunofluorescence staining of air-liquid interface (ALI) cultures representative of two donors (P1 and P2). Cultures were plated three passages after transduction with *TP63-*shRNA or EV control vectors. Staining shows the presence of mucosecretory and multiciliated cells (MUC5AC, gray; ACT, magenta). Scale bars = 100 µm. **F)** Representative heatmaps from analysis of time-lapse images of ALI cultures derived from P basal cells transduced with *TP63*-shRNA (lower) or EV (upper). Cilia beat frequency is indicated by color. **G)** Coverage by motile cilia in 3-6 fields of view in each of 1-2 replicate cultures (paired t-tests, n = 6 donors, indicated by shape).

After four passages of *TP63* knockdown, cells were plated in ALI conditions to assess the effect of knockdown on differentiation. Empty vector–transduced cells consistently formed epithelial layers with intact barrier function (TEER; 6/6; **Supplementary Fig. 7D**). Following TP63 knockdown, 3/6 donors showed comparable TEER, 2/6 showed reduced TEER within the lower normal range, and 1/6 failed to form a barrier (**Supplementary Fig. 7D**). Whole-mount immunofluorescence staining of ciliated (ACT) and mucosecretory (MUC5AC) cell types in ALI cultures showed reduced multiciliated cell differentiation after *TP63* knockdown (**Fig. 4E**). This was consistent with reduced coverage by motile cilia in ALI cultures derived from basal cells transduced with *TP63* shRNA, identified by high-speed time-lapse imaging and quantification with an ImageJ macro (CiliaFreqMap^51^; **Fig. 4F-G**).

Bulk RNA sequencing of pediatric cells after four passages of *TP63* knockdown revealed 1445 significant differentially expressed genes, 642 of which showed strong p63-dependent expression with at least a 1.5-fold change (375 up- and 267 downregulated; **Fig. 5A**). To further characterize the p63-dependent transcriptome in airway basal cells, differential expression of genes was compared between the *TP63* knockdown and overexpression RNA sequencing datasets (**Fig. 5B**). 1,301 genes were significantly differentially expressed in at least one of the datasets. Of these, the expression of 1,151 genes (88.4%) trended in opposite directions following *TP63* knockdown or overexpression (e.g. downregulated after knockdown and upregulated after overexpression), with 520 (40%) having at least 1.2-fold change in both datasets (**Fig. 5B**). Comparing this set of p63-dependent genes with the published meta-analysis of p63 expression modulation studies^50^ showed that 1,037 (90%) of the genes identified had previously been reported to display p63-dependent expression in at least one study. To assess how the changes induced by *TP63* knockdown related to those seen during aging (**Fig. 2A-D**), significant pathways identified by GSEA analysis with MSigDB hallmark gene sets were compared between aging and knockdown RNA sequencing datasets (**Fig. 5C**). Of 27 pathways identified across both datasets, 25 fell within two clusters, one representing pathways enriched in pediatric cells and pediatric cells transduced with the empty vector shRNA control, and the second representing pathways enriched in older adult cells and pediatric cells transduced with *TP63* shRNA (**Fig. 5C**). Ten of these pathways were identified in both datasets, with 15 pathways identified in only one dataset (**Fig. 5C**). This suggested that the reduction of p63 expression in pediatric cells was driving a molecular phenotype similar to that observed in older adult cells.

**Figure 5:**
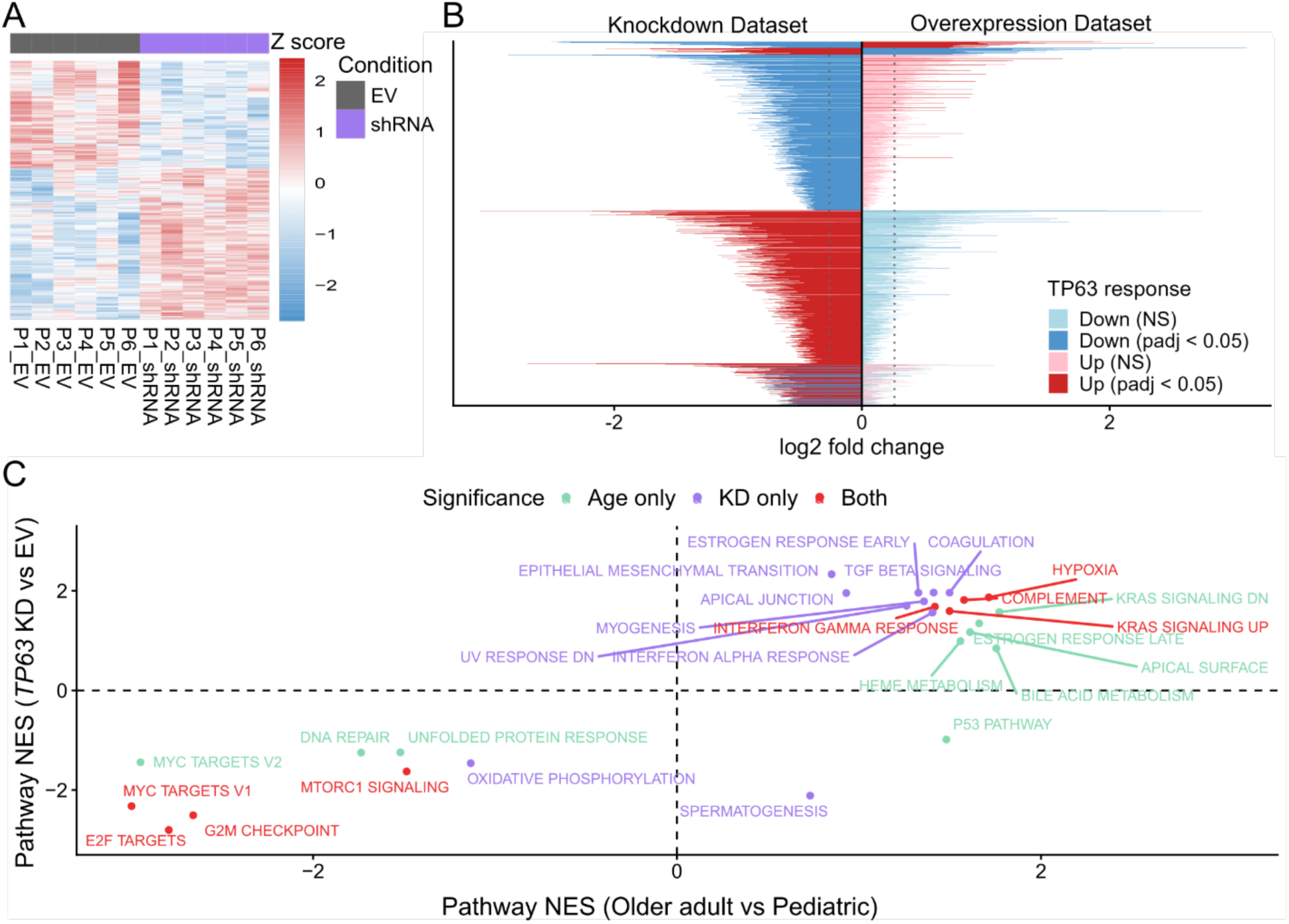
TP63 knockdown induces the enrichment of aging-associated pathways in pediatric cells. **A)** Differentially expressed genes (log2 fold change > 0.58 and adjusted p-value < 0.05) between pediatric (P) donors four passages after transduction with either *TP63*-shRNA or empty vector (EV). Normalized counts were log2-transformed and z-score scaled by gene for visualization. Sample order was fixed. Colored bar (top) indicates condition. **B)** Comparison of log2 fold changes of genes between *TP63* knockdown (left) or overexpression (right) RNA sequencing datasets. Genes significantly differentially expressed (adjusted p-value < 0.05) in at least one of the datasets were included. Direction of differential expression and significance are indicated by color. Genes are clustered by pattern of expression. Dashed lines indicate a log2 fold change of 0.263 (1.2-fold change in expression). **C)** Gene set enrichment analysis was performed using hallmark gene sets on genes ranked by log2 fold change from differential expression analysis. Concordance of pathway enrichment between aging and *TP63* knockdown RNA sequencing datasets visualized with a scatter plot comparing normalized enrichment scores (NES) of all pathways significantly enriched (adjusted p-value < 0.05) in at least one of the datasets (indicated by point color). Positive NES on the x axis denotes enrichment within older adult (OA) compared to P; positive NES on the y axis denotes enrichment within *TP63* knockdown compared to EV.

## Discussion

This study investigated the functional and molecular consequences of aging in primary human airway basal cells, using nasal brush biopsy-derived cultures from pediatric and older adult donors. We observed reduced clonogenicity and proliferative longevity in older adult compared to pediatric basal cells, that these functional differences are associated with a broad transcriptomic signature enriched for p63-dependent gene expression, and that p63 itself is downregulated in older adult cells. Lentiviral knockdown or overexpression studies show that p63 expression levels affect basal stem cell function, and integrative transcriptomic analysis defines the p63-regulated gene network in primary human airway basal cells.

Expansion of nasal basal cells from limited starting material in the majority of both pediatric (28/29) and older adult (8/12) donors demonstrates the feasibility of *in vitro* modeling of the aged airway epithelium in recently described cell culture conditions supplementing medium with the small molecule inhibitors Y-27632 and WS6^43^. The lower rate of successful culture initiation from older adult donors may reflect a reduced proportion of basal cells capable of *ex vivo* expansion in older adults, or reduced resilience of aged basal cells to the stresses of isolation and expansion.

Age-dependent, epithelial cell-intrinsic functional differences have been reported in basal cells, with lower clonogenic potential in adult cells compared to pediatric cells^3,8^ and higher replication of SARS-CoV-2 in ALI cultures derived from older adult donors^6^. In culture, basal cells are a heterogeneous population of cells with variable clonogenic potential^52^, and our data from both colony formation assays and organoid cultures suggest that pediatric cultures contain a greater proportion of clonogenic cells from the outset. The colony-forming cell frequency of both pediatric and older adult cell cultures decline during culture, but older adult cell cultures reach replicative limits earlier. This is consistent with clonal conversion to more differentiated transient amplifying progenitors, both during aging and culture^53^.

We also found substantial age-associated transcriptomic changes: early passage cultured basal cells from children were enriched with transcripts associated with cell proliferation and DNA repair, consistent with their higher proliferative potential, while culture condition-matched^54^ older adult basal cells were enriched for transcripts associated with hypoxia and interferon gamma responses, suggesting a stressed cell state. These differences likely reflect a combination of cell-intrinsic aging programs and the accumulated effect of lifetime environmental exposures retained through epigenetic memory in cultured basal cells^55,56^.

Analysis of key airway basal cell transcription factors showed downregulation of *SOX2* and *TP63* in cultured basal cells with aging. While neither protein has been linked to epithelial aging, SOX2 is under investigation, as a reprogramming factor, in efforts to rejuvenate tissue using cellular reprogramming approaches^57^. Lower protein expression of p63 in early passage older adult basal cells, as well as enrichment for p63 target genes within those differentially expressed between pediatric and older adult basal cells, led us to investigate p63’s role further. Unsupervised, hierarchical clustering on p63-dependent gene expression separated samples not by donor or age group, but by function. The two pediatric and two older adult donors with the lowest clonogenic potential clustered together with the remaining older adult samples, while the four pediatric donors with the highest colony forming efficiency formed a distinct cluster. This indicates that the p63-associated transcriptomic signature tracks stem cell functionality rather than chronological age alone. This supports prior data from studies which demonstrate that cells expressing high levels of p63 protein are those capable of long-term tissue regeneration following transplantation^11^ and suggests that low p63 activity is a shared feature of functionally depleted basal cells regardless of donor age.

Doxycycline-induced overexpression of *ΔNp63α* in older adult basal cells led to a modest but statistically significant increase in total population doublings, with cultures overexpressing *ΔNp63α* proliferating for one to two passages longer than paired controls. *ΔNp63α* overexpression improved the maintenance of colony forming efficiency across serial passages. This further supports the observation that p63-regulated differentially expressed genes in cultured pediatric versus older adult basal cells clustered by function rather than donor age.

Conversely, constitutive shRNA knockdown of *TP63* in pediatric basal cells recapitulated aspects of the aged phenotype: a significant reduction in cumulative population doublings, progressive decline in colony forming efficiency across passages, and in the most severely affected donors, failure to proliferate beyond the seventh passage following transduction. The consistent reduction in colony forming efficiency across all six donors at multiple passages, combined with the observation that residual colonies in knockdown cultures arose from GFP-negative, *TP63*-expressing cells, further supports that p63 maintains basal cell clonogenicity. Differentiation of shRNA-transduced cultures was also affected, with *TP63* knockdown leading to reduced production of multiciliated cells. These data demonstrate a key functional role for p63 in colony forming potential, long-term culture and differentiation of primary airway basal cells, extending prior evidence from immortalized airway^16,17^, cancer cell lines^17^, and primary murine tracheal basal cells^18^.

Similar to a previous study in primary murine mammary epithelial cells, where *ΔNp63* overexpression improved mammosphere colony forming efficiency and serial passaging^30^, we observed functional benefits to overexpressing ΔNp63α in older adult cells. However, it is notable that overexpression was insufficient to reverse age-associated functional decline. The broader transcriptional and epigenetic landscape of older adult cells has been shaped by years of environmental exposure, progressive epigenetic drift, and the accumulation of age-associated chromatin changes. These may constrain the transcriptional response to p63 restoration even when p63 occupancy at its target sites is re-established. p63 function is known to be strongly dependent on chromatin context and co-factor availability^24,36,58^. Additionally, since low levels of *ΔNp63β* and *ΔNp63ε* expression were detected, contributions from other p63 isoforms cannot be excluded.

The mechanistic basis for p63 protein decline with age in airway basal cells remains to be established. Protein expression decline was more consistently detected than RNA expression changes in our study, so post-transcriptional regulation, such as miRNA targeting of *TP63* mRNA^59^ or changes in proteasomal degradation via phosphorylation-dependent ubiquitination^60^, may be relevant. Upstream, age-associated changes to the ECM and to integrin-mediated adhesion signaling could provide an environmental driver of *TP63* downregulation, given established links between hemidesmosomal signaling, YAP pathway activity, and p63 expression in stratified epithelia^14,61,62^. Since p63 itself regulates integrin expression^63,64^, a self-reinforcing feedback loop between ECM remodeling and declining p63 expression is also plausible.

Based on our data, we propose a model in which p63 acts as a master regulator of airway basal cell clonogenicity and self-renewal whose declining abundance with age drives progressive impairment of epithelial function. In aged epithelium, reduced p63 activity pushes cells toward a stressed state that is prone to clonal conversion and therefore exhausts stem/progenitor cell populations and drives epithelial dysfunction. We anticipate that further dissection of the upstream regulators and downstream effectors of p63 in the airway will reveal new mechanisms of epithelial aging and identify targets for strategies aiming to preserve regenerative capacity in the aging lung.

## Materials and Methods

### Basal cell isolation from nasal brushings

Nasal brushings were taken from regions of normal nasal airway in non-smokers with no current airway disease (**Supplementary Table 1**). Tissues were transferred to the laboratory on ice in transport medium (αMEM (Gibco, 22571020) supplemented with 100 U/mL penicillin, and 100 μg/mL streptomycin, 10 μg/mL gentamicin (Gibco, 15710-049), and 250 ng/mL amphotericin B (Fisher, 10346503). Tissues were stored at 4°C overnight. Brushes were then incubated in 0.1% trypsin-EDTA (Sigma-Aldrich, 59418C) in RPMI medium (Gibco, 21875-034) for 20 minutes at 37°C with vigorous shaking every two minutes, followed by treatment with red blood cell lysis buffer (Life Technologies, 00-4333-57) for 10 minutes at room temperature.

### Epithelial cell co-culture with 3T3-J2 cells

The epithelial cell growth medium was FAD+Y+WS6 (FAD containing 5 μM Y-27632 and 100 nM WS6)^43^, which is a modified ‘conditional reprogramming’ medium^65,66^. 3T3-J2 mouse embryonic fibroblasts (Kerafast) were cultured in DMEM with sodium pyruvate (Gibco, 41966) supplemented with 7% bovine serum (Hyclone, SH30072.04; lot AE29427271), 100 U/mL penicillin, and 100 μg/mL streptomycin (Sigma-Aldrich, P0781-100ML). 3T3-J2 cell culture, preparation of mitotically inactivated 3T3-J2 feeder layers and co-culture with nasal epithelial cells were performed as previously described^43^.

### Nasal organoid cultures

Nasal organoid differentiation was performed as previously described^67^. Briefly, medium consisted of 50% DMEM (Gibco, 41966) and 50% BEBM, supplemented with BEGM singlequot supplements (excluding amphotericin B, triiodothyronine, and retinoic acid; Lonza CC-3171 and CC-4175). All-trans retinoic acid (100 nM; Sigma-Aldrich, R2625, stock 10 mM in ethanol) was freshly added before use. 5 μM Y-27632 was added to the organoid medium for cell seeding only. Ultra-low adhesion 96-well plates (Corning) were coated with 30 µL 25% Matrigel (Corning, 354230; growth factor-reduced) in organoid medium and incubated at 37°C for 30 minutes to set. Epithelial cells were seeded at 2,500 cells/well in 65 µL 5% Matrigel in organoid medium (supplemented with 5 µM Y-27632 and 100 nM retinoic acid). Organoids were incubated at 37°C for 21 days, with 50 µL fresh organoid medium (supplemented with 100 nM retinoic acid) on days 3, 10 and 17. At day 21, whole-well, phase-contrast, tile-scan images were taken for quantification of organoid number and size using OrgaQuant^68^. Subsequently, organoids were collected in ice-cold PBS, with each well of the same condition pooled together. Organoids were fixed in 4% paraformaldehyde (PFA) on ice for 30 minutes and resuspended in HistoGel^TM^ (Epredia, HG4000-012). HistoGel^TM^ plugs were transferred to 70% ethanol and kept at 4°C until proceeding with paraffin processing.

### Air-liquid interface (ALI) cultures

Semi-permeable membrane inserts (0.4 µm pore size, 0.33 cm^2^ growth area; Corning, 3470 or Sarstedt, 83.3932.041) were coated with 50 µg/mL rat tail collagen I (Corning, 354236) in 0.02 N acetic acid (Sigma-Aldrich, 1.00063.1011) for one hour at room temperature. Airway basal cells were plated onto the collagen-coated membrane supports (2-3 × 10⁵ cells in 250 µL epithelial cell growth medium). An additional 500 µL of epithelial growth medium was added to the basolateral chamber. After 24 hours, the apical medium was removed and the basolateral compartment was supplied with PneumaCult-ALI medium (STEMCELL Technologies, 05001, prepared as per manufacturer’s instructions) containing 20 U/mL penicillin and 20 μg/mL streptomycin, 4 µg/mL heparin (Sigma-Aldrich, H4784) and 0.48 µg/mL hydrocortisone (Sigma-Aldrich, H0888). Cultures were fed three times per week. Mucus was cleared by gentle apical washing with PBS once per week. ALI cultures were maintained for 28-35 days in a 37°C, 5% CO_2_, high humidity incubator prior to downstream analyses.

Transepithelial electrical resistance (TEER) was measured using an EVOM manual resistance meter (World Precision Instruments). For top-down ciliary beat frequency (CBF) measurements, high-speed time-lapse recordings were acquired at 37°C with 5% CO₂. Ciliary beat frequency and active ciliated area were quantified using the CiliaFreqMap ImageJ macro^51^. A corresponding static GFP fluorescence image was captured for each field of view.

### Tissue processing, embedding and sectioning

Formalin-fixed organoids and ALI cultures embedded in HistoGel^TM^ were processed in a Leica TP1050 processor and embedded in type 6 paraffin wax (Epredia, 8336) using an embedding station (Sakura Tissue-TEK TEC). Samples were sectioned at 5 µm thickness using a Microm HM 325 microtome and stored at 4°C until use.

### Immunofluorescence staining and imaging

For immunofluorescence staining of organoid sections, slides were de-waxed using an automated protocol (Sakura Tissue-Tek DRS). Samples were blocked at room temperature for one hour in 1% bovine serum albumin (BSA; Merck, 1.12018.0100), 5% normal goat serum (NGS; Gibco, 16120064), and 0.1% Triton X-100 (Sigma-Aldrich, X-100) in PBS. Primary antibodies (**Supplementary Table 2**) were diluted in blocking buffer and incubated on samples overnight at 4°C. Secondary antibodies (**Supplementary Table 2**) were diluted 1:1000 in 5% NGS and 0.1% Triton X-100 and incubated on samples at room temperature for three hours in the dark. Samples were then stained for 20 minutes in DAPI (Sigma-Aldrich, D9542) diluted 1:5,000 in PBS from a 1 mg/mL stock. Organoid sections were covered with a coverslip with Immu-Mount (Thermo Fisher Scientific, 9990402).

Whole-mount ALI cultures were transferred to 1.5 mL Eppendorf tubes and blocked in PBS containing 2% BSA, 5% NGS, and 0.1% Triton X-100 for one hour at room temperature at 20 rpm on a see-saw rocking platform. Primary antibodies (**Supplementary Table 2**) were diluted in blocking buffer and incubated overnight at 4°C at 60 rpm on a mini orbital shaker (Stuart, SSM1). Secondary antibodies (**Supplementary Table 2**) were diluted 1:1000 in 2% BSA, 5% NGS, 0.1% Triton X-100 in PBS and applied for three hours at room temperature in the dark on a see-saw rocking platform. DAPI (Sigma-Aldrich, D9542) was applied at 100 ng/mL for 20 min in the dark on a see-saw rocking platform. Inserts were mounted with Immu-Mount within imaging spacers (Grace Bio-Labs, 654006).

### Colony formation assays

Epithelial basal cells were seeded at 1,000 cells per well in three wells of a six-well plate prepared with 3T3-J2 feeder layers the day before. After 7 days, plates were fixed in 4% PFA for five minutes at room temperature and stained with 1% crystal violet solution (Sigma-Aldrich) for a further five minutes. Colonies (≥ 10 cells) were manually counted under a brightfield microscope. Colony forming efficiency was calculated as: (number of colonies / number of seeded cells) x 100. Plates were imaged on an Epson Perfection V600 flatbed scanner.

### Population doublings

Population doublings were calculated as PD = 3.32 × [log(cells harvested/cells seeded)].

### Lentiviral production

To generate stocks of lentiviral transfer plasmids, 10-β competent *E. coli* (NEB, C3019H) were transformed according to the manufacturer’s protocol. Plasmids were prepared from transformed bacteria with Qiagen HiSpeed maxi-prep kits (Qiagen, 12662).

Lentivirus was produced by transfection of HEK293T cells that were cultured in DMEM supplemented with 10% FBS and 100 U/mL penicillin, and 100 μg/mL streptomycin, in incubators at 37°C, 5% CO_2_. To generate viral supernatants, HEK293T cells were co-transfected at 50% confluency in a T175 with 20 µg transfer plasmid, 7 µg envelope plasmid pMD2.G (a gift from Didier Trono; Addgene plasmid 12259) and 13 µg packaging pCMV-dR8.74 (a gift from Didier Trono; Addgene plasmid 22036). Transfections were performed with JetPEI transfection reagent (Polyplus Transfection, 101000053) following the manufacturer’s protocol. Transfected cells were incubated at 37°C and viral supernatants were collected after 48 hours and concentrated with PEGit viral precipitation solution (System Biosciences, LV810A-1). The lentiviral pellet was resuspended in 1/100th of the original supernatant volume in ice-cold PBS, aliquoted and stored at −80°C. Viral aliquots were single-use.

Viral titration was performed by transduction of HEK293T cells with serial dilutions (1:10; 0.03-4 µL/mL) of lentiviral particles prepared in DMEM + 10% FBS containing 4 µg/mL polybrene. Cells were incubated at 37 °C, 5% CO₂ for 7 hours before medium was changed to remove viral particles. 72 hours after transduction, cells were analyzed by flow cytometry. Viral titer was calculated using fluorophore-positive populations within the linear range (10–20% positivity).

Titer (TU/ml) = (N x P) / (V x D), where:

TU = transduction unit, N = number of cells in one well on Day 1, P = percentage of positive cells, V = virus volume used in the first well in mL (0.004 in this protocol), D = dilution fold (1 for 4 µL well, 0.5 for 2 µL well, 0.25 for 1 µL well, etc.).

### Lentiviral transduction of primary epithelial cells

150,000 cells were transduced in one well of a 6-well plate (equivalent to 15,000-16,000 cells/cm^2^) in 1.5 mL sterile-filtered (0.22 µm; SLS, B2B06412) EpMED that had been collected from proliferating epithelial cells (50-90% confluency) for 24 h and supplemented with 10 ng/mL EGF, 5 μM Y-27632 and 100 nM WS6 (conditioned medium). Conditioned medium containing polybrene (8 µg/mL final concentration) and lentiviral viral particles equivalent to MOI = 1 (Volume of virus (mL) = MOI x (number of cells) / Titer (TU/mL)). Cells were centrifuged at 920 × g for one hour at 30°C and then incubated at 37°C, 5% CO_2_ for 5-7 hours. Medium was refreshed and 3T3-J2 fibroblast feeder cells were added at 20,000/cm^2^. Cells were expanded for 3-4 days before selection by fluorescence activated cell sorting (FACS). Cells transduced with both the Tet-On and *ΔNp63α*-mCherry vectors (**Supplementary Table 3**) required selection with 250 µg/mL hygromycin B for four days prior to FACS.

### RNA and protein extraction

Cells were cultured to 80% confluence and growth medium was refreshed the day before RNA or protein collection. 3T3-J2 feeder cells were removed by a trypsin/EDTA-wash cycle immediately before collection and cells were then washed with 1 ml epithelial cell culture medium, followed by 1 ml PBS on ice. RNA lysates were collected by scraping cells using plastic plate scrapers (VWR, 734-2602) into 600 µL Lysis buffer from the PureLink RNA Mini Kit (Invitrogen, 12183018A) supplemented with 10 µL β-mercaptoethanol (Serva, 39563.02), or 350 µL RLT buffer (Qiagen, 80204). RNA lysates were stored at −80°C until processing, when they were thawed on ice and processed according to the manufacturer’s protocol (PureLink RNA Mini Kit or Qiagen All-Prep DNA/RNA Mini Kit). RNA was stored at −80°C prior to use.

Protein lysates were collected by scraping using plastic plate scrapers in 150 µL RIPA buffer (Thermo Scientific, 89900) supplemented with protease and phosphatase inhibitor cocktail (Fisher Scientific, 10025743). To remove cell debris, protein lysates were centrifuged at 14,000 x g for 15 minutes, with the supernatant transferred to a new Eppendorf. Protein was stored at - 80°C prior to use.

### Quantitative real-time polymerase chain reaction (RT-qPCR)

RNA was quantified using a DeNOVIX DS-11FX series spectrophotometer. 1 µg was reverse transcribed to cDNA using qScript cDNA SuperMix (VWR, 95048-100) following the manufacturer’s protocol. RT-qPCR was performed with Power SYBR Green PCR Master Mix (Thermo, 4368706), and 5 µM forward and reverse primer mixes (Supplementary Table 4). Plates were run in a QuantStudio™ 5 Real-Time PCR System (Applied Biosystems). RNA expression was quantified by the Delta Ct method, relative to two housekeeping genes: glyceraldehyde 3-phosphate dehydrogenase (*GAPDH*) and ribosomal protein S13 (*RPS13*).

### Bulk and single-cell RNA sequencing analysis

Bulk RNA sequencing was performed by Novogene. Library preparation was performed using the NEBNext® Ultra™ RNA Library Prep Kit to construct cDNA libraries with an insert size of approximately 250-300 bp. Libraries were sequenced on the Novaseq X Plus platform using a PE150 (paired-end 150 bp) sequencing strategy.

Data were processed using the nf-core/rnaseq pipeline (v3.22.2^69^) with Nextflow (v25.10.0 build 10289^70^). Raw sequencing reads were assessed for quality using FastQC and aggregated with MultiQC. Adapter trimming and quality filtering were performed by Trim Galore. Reads were aligned to the human reference genome (GRCh38) by STAR using the Ensembl v110 annotation. Transcript abundance was quantified by Salmon. The pipeline was executed using default parameters using an Apptainer container environment. Salmon quantification files were used for downstream analysis and visualization in RStudio (v4.4.2).

Transcript-level quantifications generated by the pipeline were summarized to gene-level counts using tximport (v1.34.0^71^). Differential expression analysis was performed using DESeq2 (v1.46.0^72^). Genes located on autosomal chromosomes (1-22) were retained for analysis. Additionally, only genes with a minimum of 50 counts in all samples of one experimental group were retained for analysis. For *TP63* knockdown and overexpression datasets, models included donor as a covariate to account for inter-individual variability. Log2 fold changes were shrunk using the apeglm (v1.28.0^73^) method. Adjusted p-values were calculated using the Benjamini– Hochberg procedure, with significance defined as padj < 0.05. Gene annotations were obtained using org.Hs.eg.db (v3.20.0^74^) via Annotation Dbi (v1.68.0^75^) and biomaRt (v2.62.1^76^).

Normalized single-cell RNA sequencing data of *in vivo* airway samples collated from seven studies^77–83^ was downloaded in H5AD format and imported into RStudio using zellkonverter (v1.16.0^84^) and SingleCellExperiment (v1.28.1^85^) frameworks. Data were converted into a Seurat (v5.4.0^86^) object for downstream analysis. Cells were subset to include only data from healthy pediatric (0-18 years) and older adult (51-90 years) donors. Pre-computed UMAP embeddings were incorporated directly for visualization. Cell type annotations provided in the metadata were used to label clusters. Differential expression analysis of *TP63* and *SOX2* within basal cells expressing the respective genes was performed using Wilcoxon rank-sum tests within Seurat.

### Western blotting

Protein concentration was quantified using a Pierce BCA Protein Assay kit (Thermo Scientific, 23225), following the manufacturer’s microplate protocol. Western blots were performed with 10 µg of protein per sample. Protein samples were thawed on ice and denatured in 2x Laemmli buffer (Sigma-Aldrich, S3401-1VL) with a 10 minute incubation at 95°C. Denatured protein samples and PageRuler Prestained 10-180 kDa protein ladder (Thermo Fisher, 26616) were loaded into 4-12% Bis-Tris buffer gels (Thermo Scientific, NW04125BOX) and ran with MOPS SDS running buffer (Thermo Fisher, B0001) at 120 V for approximately 75 minutes.

Protein was wet-transferred onto an Immobilon-P PVDF membrane (Millipore, IPVH00010) for two hours at 70 V at 4°C in wet transfer buffer (25 mM Tris, 192 mM glycine, 20% methanol in dH_2_O). Membranes were blocked with 5% milk powder (VWR Chemicals, 84615.0500) in PBS with 0.1% TWEEN20 (PBST) for one hour at room temperature. Membranes were incubated with primary antibodies (**Supplementary Table 5**) diluted in blocking buffer at 4°C overnight with rotation. Membranes were incubated with HRP-conjugated anti-rabbit secondary antibody (**Supplementary Table 5**) diluted in blocking buffer for two hours at room temperature with rotation. Membranes were covered with Crescendo HRP substrate (Millipore, WBLUR0100) for one minute before chemiluminescence imaging on a BioRad ChemiDoc MP imaging system. As a loading control, membranes were subsequently incubated with HRP-conjugated anti-α-Tubulin antibody (**Supplementary Table 5**) diluted in blocking buffer for one hour at room temperature, with rotation. Relative band intensity was quantified in ImageJ.

## Data and code availability

Basal cell RNA sequencing data generated in this study has been deposited with GEO (GSE328414). Code for analysis and data visualization of RNA sequencing data are available at Zenodo (doi: 10.5281/zenodo.19736892).

## Acknowledgements

A.S.F. thanks the members of his PhD thesis committee, Dr Manuela Plate, Prof Paolo De Coppi, Dr Marc Amoyel, Dr Yang Li and Dr Joana Rodrigues Simoes Da Costa for their guidance. The authors thank Dr Maral Rouhani (UCL Division of Medicine, University College London, U.K.) and Mr Colin Butler (Great Ormond Street Hospital for Children, London, U.K.) for collecting the nasal brush biopsy samples used in this study. The authors also thank Dr Ayad Eddaoudi and Panayiota Constantinou (both Flow Cytometry Core Facility, UCL Institute of Child Health, University College London, U.K.), Dr Dale Moulding (Microscopy Core Facility, UCL Institute of Child Health, University College London, U.K.) and UCL Genomics (RRID:SCR_027010) for their training and assistance. Finally, the authors thank members of the EpiCENTR group for feedback on the draft manuscript.

## Author contributions

A.S.F. and R.E.H. conceived the study and planned experiments. A.S.F. performed cell culture experiments and bioinformatic analyses with contributions from J.C.O. (to *TP63* shRNA experiments), B.M.A. (to colony formation assays) and L.H. (to qPCR experiments). T.B. performed library preparation and sequencing experiments. A.S.F. wrote the first manuscript draft, which was revised by R.E.H. All authors provided feedback on the manuscript.

## Ethics declarations

Nasal brush biopsies were obtained from patients with informed consent from patients or their parents/guardians. Ethical approval was obtained through a National Research Ethics Committee (Living Airways Biobank, REC 24/NW/0168).

## Supplementary Figures

**Supplementary Figure 1:**
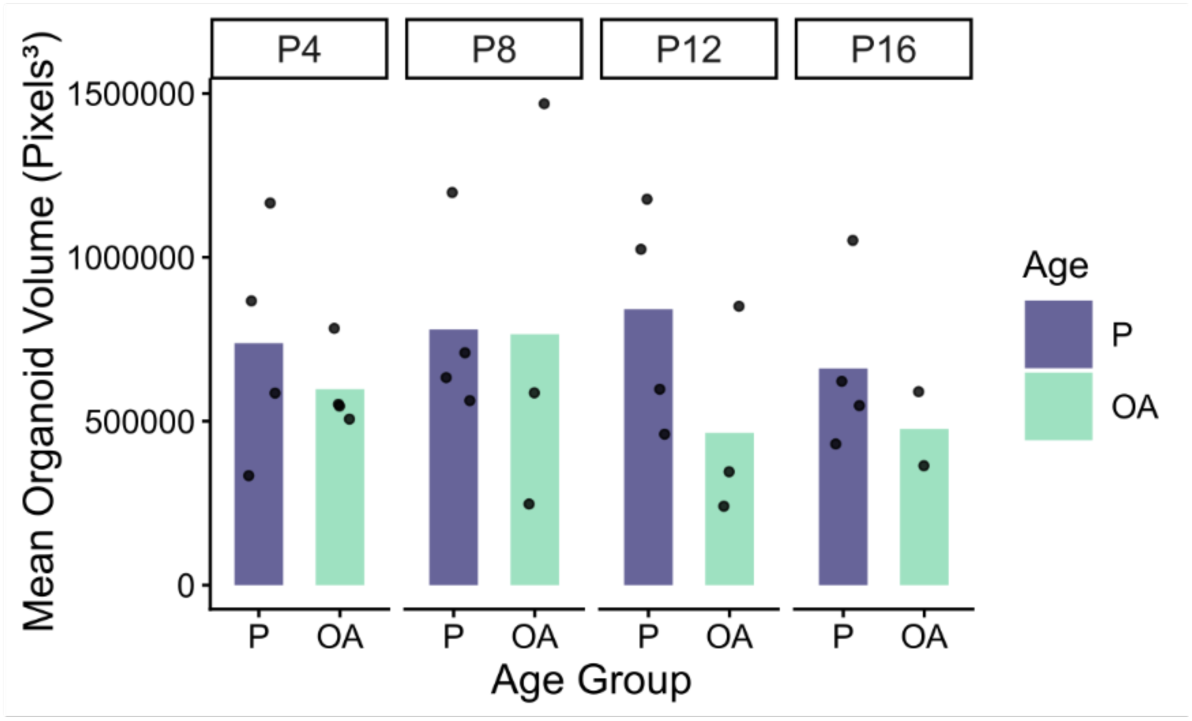
Organoids derived from pediatric and older adult basal cells are comparable in size. Organoid size from P or OA basal cells at passage 4, 8, 12 or 16. Wilcoxon signed-rank test compared mean organoid number indicated by bars, points show the mean of 8 well replicates per donor (n = 4 P, 2-4 OA donors; lack of stars indicates non-significance).

**Supplementary Figure 2:**
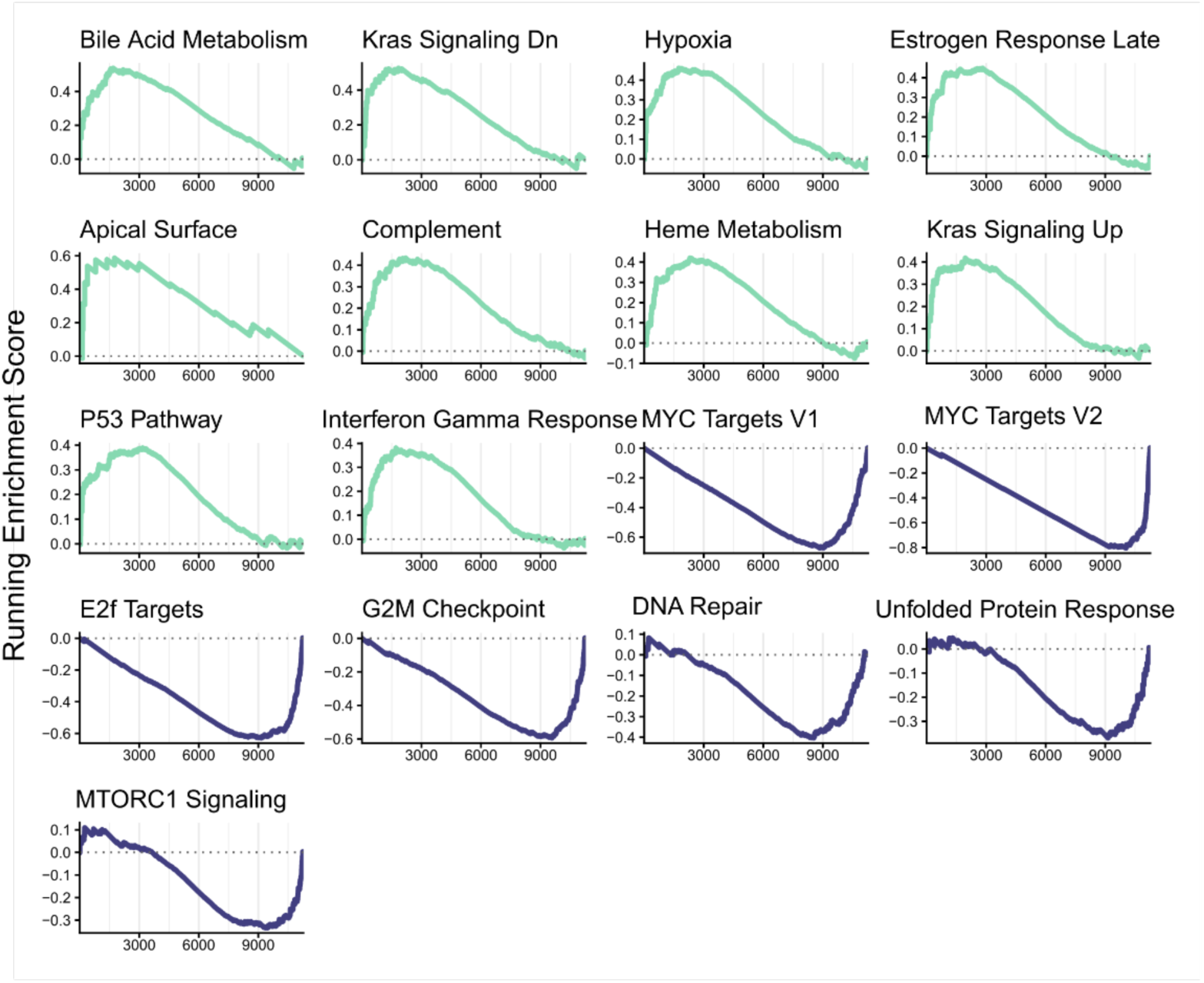
Gene set enrichment analysis of RNA sequencing data comparing cultured pediatric (P) with older adult (OA) airway basal cells. All pathways significantly enriched (p < 0.05) within P (purple) and OA (green) transcriptomes. Curves show running enrichment scores.

**Supplementary Figure 3:**
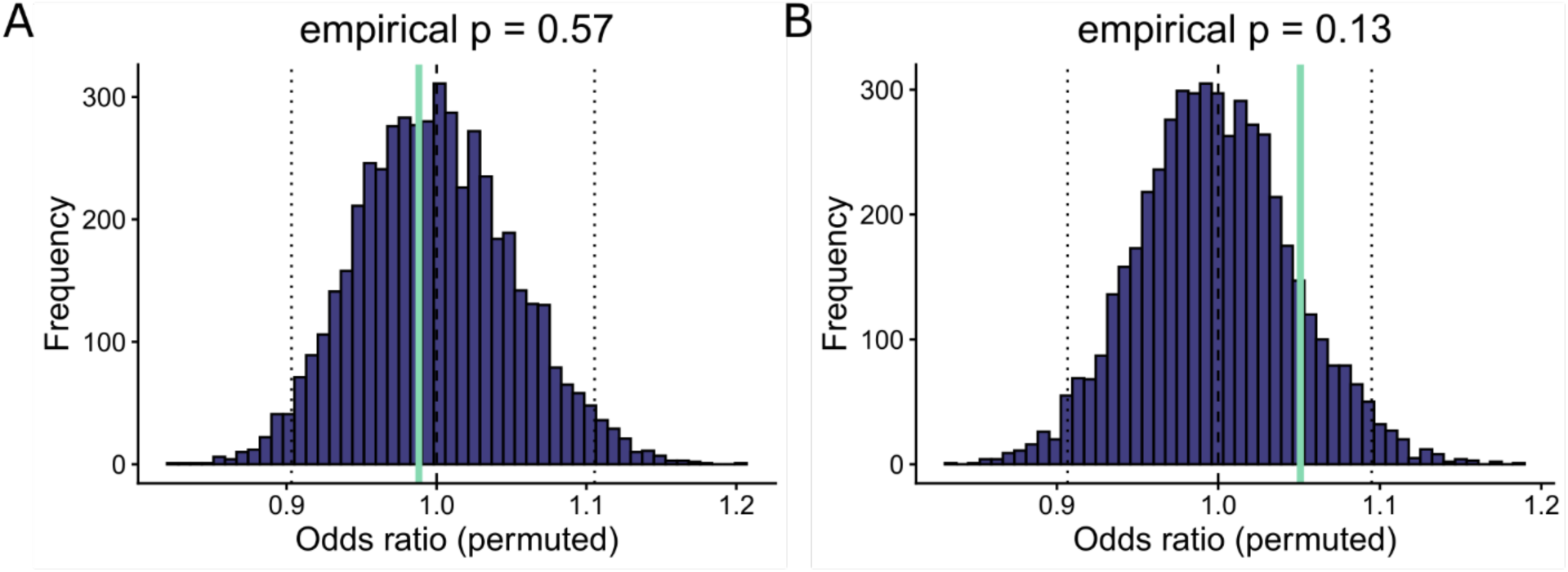
Fisher’s exact test for enrichment of SOX2- or p63-bound genes among age-associated differentially expressed genes. Enrichment of putative SOX2 **(A)** or p63 **(B)** target genes from the ChIP-X Enrichment Analysis database among differentially expressed genes (adjusted p-value < 0.05, |log2 fold change| ≥ 0.263) was assessed using a Fisher’s exact test. The histogram shows the outcome of 5,000 random permutations, with 95% confidence interval indicated by dashed lines. The observed odds ratio is indicated by the green line and the empirical p-value reflects the proportion of permutations exceeding the observed enrichment.

**Supplementary Figure 4:**
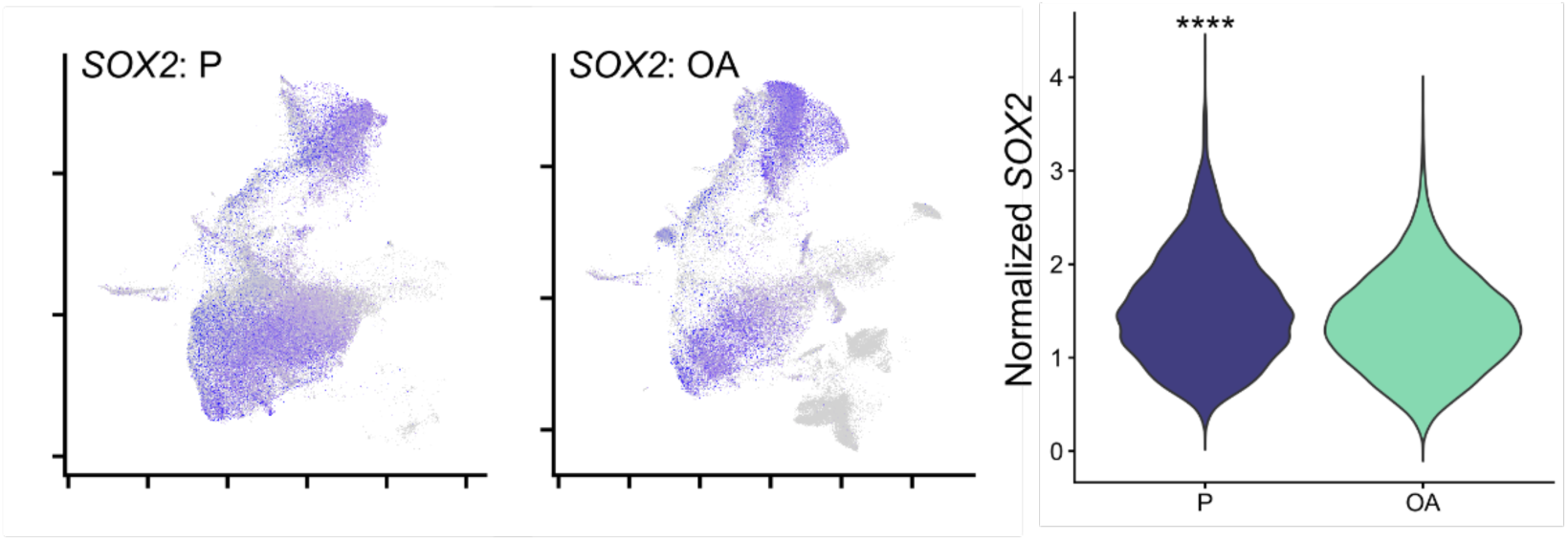
SOX2 expression in single cell RNA sequencing dataset. SOX2 expression in pediatric (P, left) and older adult (OA, middle) donors was visualized using feature plots with quantile-based cutoffs (q10–q90). Cell type annotations provided in the metadata were used to label clusters and can be found in main Figure 2H. SOX2 expression was compared in basal cells with detectable SOX2 expression (>0; Wilcoxon rank-sum test, right; P: n = 10,544 cells; OA: n = 6,336 cells).

**Supplementary Figure 5:**
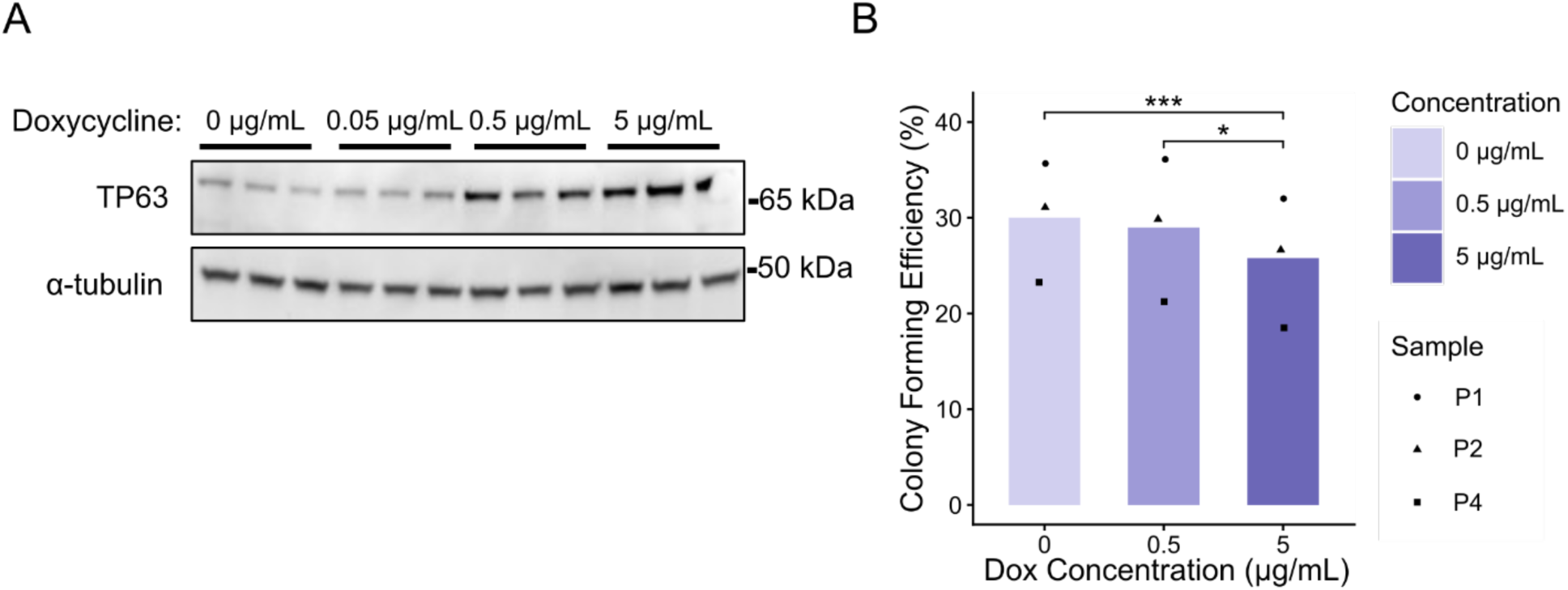
*TP63* overexpression is doxycycline concentration-dependent. **A)** Western blot showing ΔNp63α protein expression in n = 1 primary older adult donor transduced with the TP63 overexpression vectors after one passage of doxycycline induction at different concentrations (0 µg/mL, 0.05 µg/mL, 0.5 µg/mL, 5 µg/mL). α-tubulin was used as a loading control. **B)** Colony formation efficiency of primary, non-transduced pediatric basal cells. Cells were plated in medium supplemented with doxycycline (0 µg/mL, 0.05 µg/mL, 0.5 µg/mL, 5 µg/mL) for the seven-day duration of the assay. One-way ANOVA was used to test the effect of doxycycline concentration (p = 0.0009) with Tukey’s post hoc to identify significant pairwise comparisons (n = 3 donors, mean of 3 well replicates per donor).

**Supplementary Figure 6:**
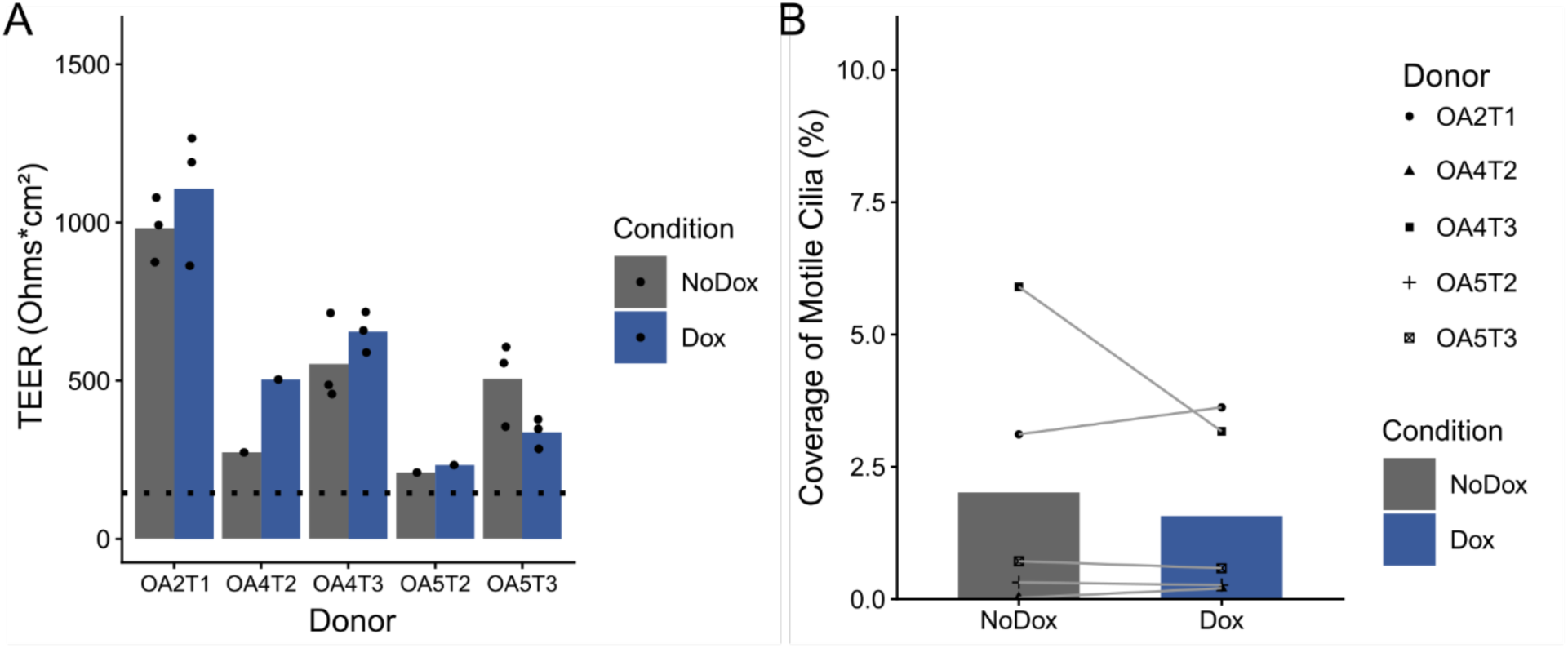
TP63 overexpression does not restore differentiation to older adult basal cells. **A)** Transepithelial electrical resistance (TEER) of air-liquid interface (ALI) cultures derived from older adult basal cells transduced with TP63 overexpression vectors and divided into +/- doxycycline conditions for three passages before plating. Dashed line marks the blank well reading (paired Wilcoxon signed-rank test; n = 4 independent transductions in 2 donor cell cultures). **B)** Coverage by motile cilia in 3-6 fields of view in each of 1-2 replicate ALI cultures. compared mean coverage (paired Wilcoxon signed-rank test; n = 5 independent transductions in 3 donor cell cultures).

**Supplementary Figure 7:**
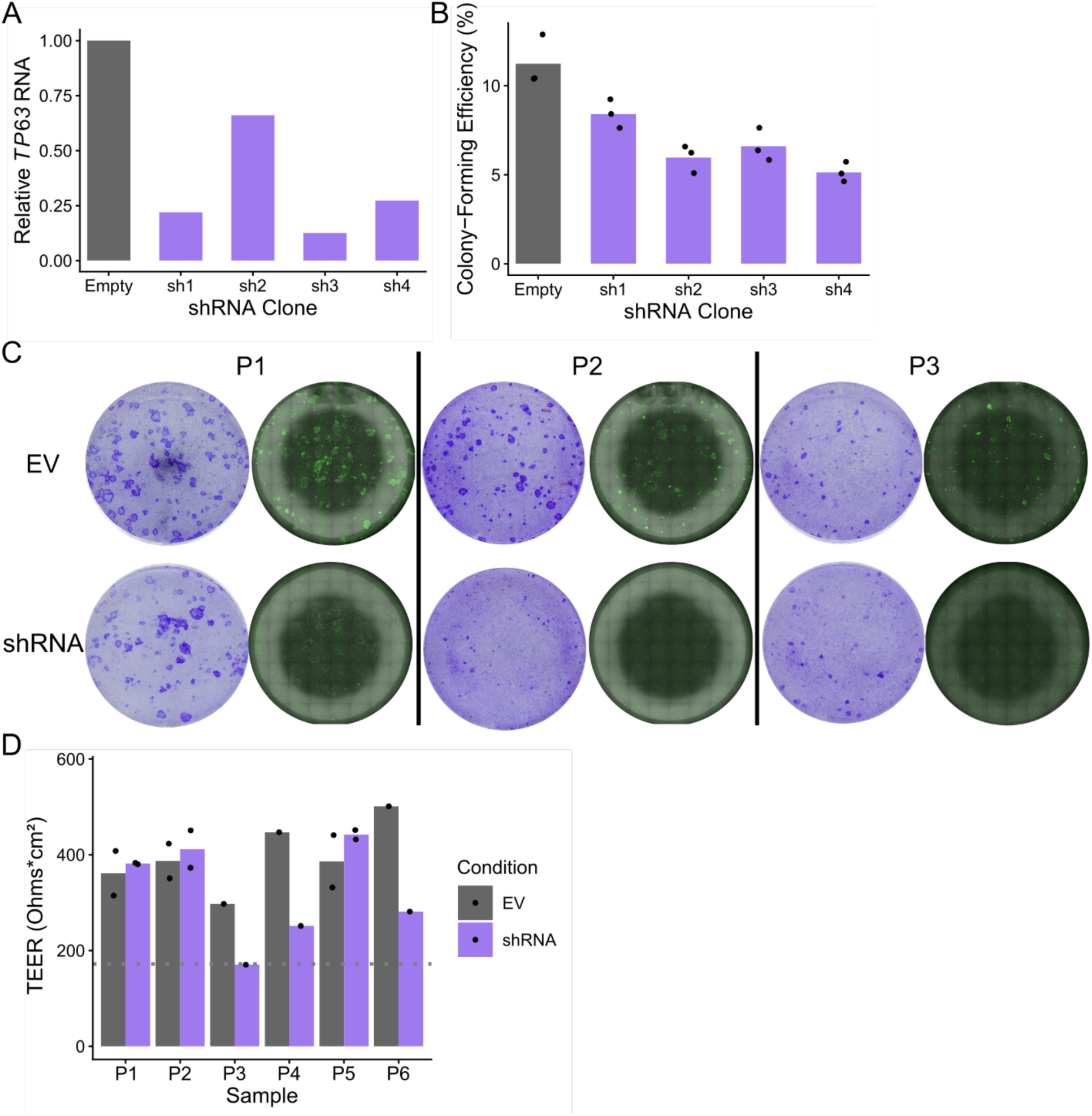
Optimization of TP63 shRNA knockdown and additional functional effects. **A)** ΔNp63 RNA expression after one passage of knockdown in n = 1 primary pediatric donor cell culture with four different TP63-shRNA clones. Expression is calculated relative to control cells transduced with an empty vector. Expression was measured using a single RT-qPCR experiment, with 3 well replicates. **B)** Colony forming efficiency of cells transduced with TP63-shRNA or empty vector (n = 1 pediatric donor). Each point represents the percentage of cells that formed a colony from a single well. Bars represent the average of 3 well replicates. **C)** Representative well images of colony formation assays plated seven passages after transduction (n = 3 donors). Flatbed scans of one crystal violet-stained colony formation assay well (left) and corresponding tile scans with merged phase-contrast and GFP channels (right) are shown. Each well is representative of 3 well replicates. **D)** Transepithelial electrical resistance (TEER) of ALI cultures derived from pediatric basal cells transduced with TP63-shRNA or empty vector. Dashed line marks the blank well reading (Wilcoxon signed-rank test; n = 6 donors).

## References

1 Miller Z, Twardowski L-M, Reader BF, Rojas M, Lehmann M, Mora AL. Mechanisms and markers of lung ageing in health and disease. Eur Respir Rev 2025; 34. doi:10.1183/16000617.0233-2024.

2 Baker JR, Beaulieu D, Avci E, Huang E, Eickelberg O, Meiners S et al. Hallmarks of the ageing lung - 10 years later. Eur Respir J 2026. doi:10.1183/13993003.01272-2025.

3 Yoshida K, Gowers KHC, Lee-Six H, Chandrasekharan DP, Coorens T, Maughan EF et al. Tobacco smoking and somatic mutations in human bronchial epithelium. Nature 2020; 578: 266–272.

4 Huang Z, Sun S, Lee M, Maslov AY, Shi M, Waldman S et al. Single-cell analysis of somatic mutations in human bronchial epithelial cells in relation to aging and smoking. Nat Genet 2022; 54: 492–498.

5 Aghali A, Koloko Ngassie ML, Pabelick CM, Prakash YS. Cellular Senescence in Aging Lungs and Diseases. Cells 2022; 11. doi:10.3390/cells11111781.

6 Woodall MNJ, Cujba A-M, Worlock KB, Case K-M, Masonou T, Yoshida M et al. Age-specific nasal epithelial responses to SARS-CoV-2 infection. Nat Microbiol 2024; 9: 1293–1311.

7 Chason KD, Jaspers I, Parker J, Sellers S, Brighton LE, Hunsucker SA et al. Age-Associated Changes in the Respiratory Epithelial Response to Influenza Infection. J Gerontol A Biol Sci Med Sci 2018; 73: 1643–1650.

8 Maughan EF, Hynds RE, Pennycuick A, Nigro E, Gowers KHC, Denais C et al. Cell-intrinsic differences between human airway epithelial cells from children and adults. iScience 2022; 25: 105409.

9 Balázs A, Millar-Büchner P, Mülleder M, Farztdinov V, Szyrwiel L, Addante A et al. Age-Related Differences in Structure and Function of Nasal Epithelial Cultures From Healthy Children and Elderly People. Front Immunol 2022; 13: 822437.

10 Rock JR, Onaitis MW, Rawlins EL, Lu Y, Clark CP, Xue Y et al. Basal cells as stem cells of the mouse trachea and human airway epithelium. Proc Natl Acad Sci U S A 2009; 106: 12771–12775.

11 Pellegrini G, Dellambra E, Golisano O, Martinelli E, Fantozzi I, Bondanza S et al. p63 identifies keratinocyte stem cells. Proc Natl Acad Sci U S A 2001; 98: 3156–3161.

12 Signoretti S, Waltregny D, Dilks J, Isaac B, Lin D, Garraway L et al. p63 is a prostate basal cell marker and is required for prostate development. Am J Pathol 2000; 157: 1769–1775.

13 Barbareschi M, Pecciarini L, Cangi MG, Macrì E, Rizzo A, Viale G et al. p63, a p53 homologue, is a selective nuclear marker of myoepithelial cells of the human breast. Am J Surg Pathol 2001; 25: 1054–1060.

14 Carroll DK, Carroll JS, Leong C-O, Cheng F, Brown M, Mills AA et al. p63 regulates an adhesion programme and cell survival in epithelial cells. Nat Cell Biol 2006; 8: 551–561.

15 Sethi I, Romano R-A, Gluck C, Smalley K, Vojtesek B, Buck MJ et al. A global analysis of the complex landscape of isoforms and regulatory networks of p63 in human cells and tissues. BMC Genomics 2015; 16: 584.

16 Arason AJ, Jonsdottir HR, Halldorsson S, Benediktsdottir BE, Bergthorsson JT, Ingthorsson S et al. deltaNp63 has a role in maintaining epithelial integrity in airway epithelium. PLoS One 2014; 9: e88683.

17 Warner SMB, Hackett T-L, Shaheen F, Hallstrand TS, Kicic A, Stick SM et al. Transcription factor p63 regulates key genes and wound repair in human airway epithelial basal cells. Am J Respir Cell Mol Biol 2013; 49: 978–988.

18 Napoli M, Wu SJ, Gore BL, Abbas HA, Lee K, Checker R et al. ΔNp63 regulates a common landscape of enhancer associated genes in non-small cell lung cancer. Nat Commun 2022; 13: 614.

19 Li Y, Giovannini S, Wang T, Fang J, Li P, Shao C et al. P63: A crucial player in epithelial stemness regulation. Oncogene 2023; 42: 3371–3384.

20 LeBoeuf M, Terrell A, Trivedi S, Sinha S, Epstein JA, Olson EN et al. Hdac1 and Hdac2 act redundantly to control p63 and p53 functions in epidermal progenitor cells. Dev Cell 2010; 19: 807– 818.

21 Truong AB, Kretz M, Ridky TW, Kimmel R, Khavari PA. p63 regulates proliferation and differentiation of developmentally mature keratinocytes. Genes Dev 2006; 20: 3185–3197.

22 Wu N, Rollin J, Masse I, Lamartine J, Gidrol X. p63 regulates human keratinocyte proliferation via MYC-regulated gene network and differentiation commitment through cell adhesion-related gene network. J Biol Chem 2012; 287: 5627–5638.

23 Yalcin-Ozuysal O, Fiche M, Guitierrez M, Wagner K-U, Raffoul W, Brisken C. Antagonistic roles of Notch and p63 in controlling mammary epithelial cell fates. Cell Death Differ 2010; 17: 1600–1612.

24 Pattison JM, Melo SP, Piekos SN, Torkelson JL, Bashkirova E, Mumbach MR et al. Retinoic acid and BMP4 cooperate with p63 to alter chromatin dynamics during surface epithelial commitment. Nat Genet 2018; 50: 1658–1665.

25 Zhang Y, Karagiannis D, Liu H, Lin M, Fang Y, Jiang M et al. Epigenetic regulation of p63 blocks squamous-to-neuroendocrine transdifferentiation in esophageal development and malignancy. Sci Adv 2024; 10: eadq0479.

26 Brauweiler AM, Leung DYM, Goleva E. The Transcription Factor p63 Is a Direct Effector of IL-4- and IL-13-Mediated Repression of Keratinocyte Differentiation. J Invest Dermatol 2021; 141: 770–778.

27 Li N, Singh S, Cherukuri P, Li H, Yuan Z, Ellisen LW et al. Reciprocal intraepithelial interactions between TP63 and hedgehog signaling regulate quiescence and activation of progenitor elaboration by mammary stem cells. Stem Cells 2008; 26: 1253–1264.

28 Ning B, Tilston-Lunel AM, Simonetti J, Hicks-Berthet J, Matschulat A, Pfefferkorn R et al. Convergence of YAP/TAZ, TEAD and TP63 activity is associated with bronchial premalignant severity and progression. J Exp Clin Cancer Res 2023; 42: 116.

29 Okuyama R, Ogawa E, Nagoshi H, Yabuki M, Kurihara A, Terui T et al. p53 homologue, p51/p63, maintains the immaturity of keratinocyte stem cells by inhibiting Notch1 activity. Oncogene 2007; 26: 4478–4488.

30 Chakrabarti R, Wei Y, Hwang J, Hang X, Andres Blanco M, Choudhury A et al. ΔNp63 promotes stem cell activity in mammary gland development and basal-like breast cancer by enhancing Fzd7 expression and Wnt signalling. Nat Cell Biol 2014; 16: 1004-15, 1–13.

31 Romano R-A, Smalley K, Magraw C, Serna VA, Kurita T, Raghavan S et al. ΔNp63 knockout mice reveal its indispensable role as a master regulator of epithelial development and differentiation. Development 2012; 139: 772–782.

32 Fessing MY, Mardaryev AN, Gdula MR, Sharov AA, Sharova TY, Rapisarda V et al. p63 regulates Satb1 to control tissue-specific chromatin remodeling during development of the epidermis. J Cell Biol 2011; 194: 825–839.

33 McDade SS, Henry AE, Pivato GP, Kozarewa I, Mitsopoulos C, Fenwick K et al. Genome-wide analysis of p63 binding sites identifies AP-2 factors as co-regulators of epidermal differentiation. Nucleic Acids Res 2012; 40: 7190–7206.

34 Bao X, Rubin AJ, Qu K, Zhang J, Giresi PG, Chang HY et al. A novel ATAC-seq approach reveals lineage-specific reinforcement of the open chromatin landscape via cooperation between BAF and p63. Genome Biol 2015; 16: 284.

35 Kouwenhoven EN, van Heeringen SJ, Tena JJ, Oti M, Dutilh BE, Alonso ME et al. Genome-wide profiling of p63 DNA-binding sites identifies an element that regulates gene expression during limb development in the 7q21 SHFM1 locus. PLoS Genet 2010; 6: e1001065.

36 Kouwenhoven EN, Oti M, Niehues H, van Heeringen SJ, Schalkwijk J, Stunnenberg HG et al. Transcription factor p63 bookmarks and regulates dynamic enhancers during epidermal differentiation. EMBO Rep 2015; 16: 863–878.

37 Mehrazarin S, Chen W, Oh J-E, Liu ZX, Kang KL, Yi JK et al. The p63 Gene Is Regulated by Grainyhead-like 2 (GRHL2) through Reciprocal Feedback and Determines the Epithelial Phenotype in Human Keratinocytes. J Biol Chem 2015; 290: 19999–20008.

38 Olsen JR, Oyan AM, Rostad K, Hellem MR, Liu J, Li L et al. p63 attenuates epithelial to mesenchymal potential in an experimental prostate cell model. PLoS One 2013; 8: e62547.

39 Jiang Y, Zheng Y, Zhang Y-W, Kong S, Dong J, Wang F et al. Reciprocal inhibition between TP63 and STAT1 regulates anti-tumor immune response through interferon-γ signaling in squamous cancer. Nat Commun 2024; 15: 2484.

40 Memmi EM, Sanarico AG, Giacobbe A, Peschiaroli A, Frezza V, Cicalese A et al. p63 Sustains self-renewal of mammary cancer stem cells through regulation of Sonic Hedgehog signaling. Proc Natl Acad Sci U S A 2015; 112: 3499–3504.

41 Somerville TDD, Xu Y, Miyabayashi K, Tiriac H, Cleary CR, Maia-Silva D et al. TP63-Mediated Enhancer Reprogramming Drives the Squamous Subtype of Pancreatic Ductal Adenocarcinoma. Cell Rep 2018; 25: 1741–1755.e7.

42 Watanabe H, Ma Q, Peng S, Adelmant G, Swain D, Song W et al. SOX2 and p63 colocalize at genetic loci in squamous cell carcinomas. J Clin Invest 2014; 124: 1636–1645.

43 Orr JC, Haughey EK, Farr AS, Pearce DR, McCarthy NA, Reddy SK, et al. WS6 enables scalable ex vivo expansion and gene editing of human basal epithelial cells. bioRxiv. 2025; : 2025.09.15.675861.

44 Gao X, Bali AS, Randell SH, Hogan BLM. GRHL2 coordinates regeneration of a polarized mucociliary epithelium from basal stem cells. J Cell Biol 2015; 211: 669–682.

45 Dong Z, Wit N, Agarwal A, Reid AJ, Dubal D, Beier S et al. Hypoxia promotes airway differentiation in the human lung epithelium. Cell Stem Cell 2025; 32: 1705–1722.e9.

46 Ochieng JK, Schilders K, Kool H, Boerema-De Munck A, Buscop-Van Kempen M, Gontan C et al. Sox2 regulates the emergence of lung basal cells by directly activating the transcription of Trp63. Am J Respir Cell Mol Biol 2014; 51: 311–322.

47 Ievlev V, Jensen-Cody CC, Lynch TJ, Pai AC, Park S, Shahin W et al. Sox9 and Lef1 Regulate the Fate and Behavior of Airway Glandular Progenitors in Response to Injury. Stem Cells 2022; 40: 778– 790.

48 Daniely Y, Liao G, Dixon D, Linnoila RI, Lori A, Randell SH et al. Critical role of p63 in the development of a normal esophageal and tracheobronchial epithelium. Am J Physiol Cell Physiol 2004; 287: C171–81.

49 Lachmann A, Xu H, Krishnan J, Berger SI, Mazloom AR, Ma’ayan A. ChEA: transcription factor regulation inferred from integrating genome-wide ChIP-X experiments. Bioinformatics 2010; 26: 2438–2444.

50 Riege K, Kretzmer H, Sahm A, McDade SS, Hoffmann S, Fischer M. Dissecting the DNA binding landscape and gene regulatory network of p63 and p53. Elife 2020; 9. doi:10.7554/eLife.63266.

51 DaleMoulding. DaleMoulding/CiliaFreqMap: initial release. Zenodo, 2025 doi:10.5281/ZENODO.17107574.

52 Reynolds SD, Carraro G, Alsudayri A, Hill CL, Stack JT, Calyeca J et al. Human bronchial basal cells are a community of variants. iScience 2025; 28: 113633.

53 Barrandon Y, Green H. Three clonal types of keratinocyte with different capacities for multiplication. Proc Natl Acad Sci U S A 1987; 84: 2302–2306.

54 Murthy S, Seabold DA, Gautam LK, Caceres AM, Sease R, Calvert BA et al. Culture conditions differentially regulate the inflammatory niche and cellular phenotype of tracheobronchial basal stem cells. Am J Physiol Lung Cell Mol Physiol 2025; 328: L538–L553.

55 Ordovas-Montanes J, Dwyer DF, Nyquist SK, Buchheit KM, Vukovic M, Deb C et al. Allergic inflammatory memory in human respiratory epithelial progenitor cells. Nature 2018; 560: 649–654.

56 Goldfarbmuren KC, Jackson ND, Sajuthi SP, Dyjack N, Li KS, Rios CL et al. Dissecting the cellular specificity of smoking effects and reconstructing lineages in the human airway epithelium. Nat Commun 2020; 11: 2485.

57 Ocampo A, Reddy P, Martinez-Redondo P, Platero-Luengo A, Hatanaka F, Hishida T et al. In Vivo Amelioration of Age-Associated Hallmarks by Partial Reprogramming. Cell 2016; 167: 1719– 1733.e12.

58 Lin-Shiao E, Lan Y, Welzenbach J, Alexander KA, Zhang Z, Knapp M et al. p63 establishes epithelial enhancers at critical craniofacial development genes. Sci Adv 2019; 5: eaaw0946.

59 Candi E, Amelio I, Agostini M, Melino G. MicroRNAs and p63 in epithelial stemness. Cell Death Differ 2015; 22: 12–21.

60 Li C, Xiao Z-X. Regulation of p63 protein stability via ubiquitin-proteasome pathway. Biomed Res Int 2014; 2014: 175721.

61 Elbediwy A, Vincent-Mistiaen ZI, Spencer-Dene B, Stone RK, Boeing S, Wculek SK et al. Integrin signalling regulates YAP and TAZ to control skin homeostasis. Development 2016; 143: 1674–1687.

62 De Rosa L, Secone Seconetti A, De Santis G, Pellacani G, Hirsch T, Rothoeft T et al. Laminin 332-Dependent YAP Dysregulation Depletes Epidermal Stem Cells in Junctional Epidermolysis Bullosa. Cell Rep 2019; 27: 2036–2049.e6.

63 Kurata S-I, Okuyama T, Osada M, Watanabe T, Tomimori Y, Sato S et al. p51/p63 Controls subunit alpha3 of the major epidermis integrin anchoring the stem cells to the niche. J Biol Chem 2004; 279: 50069–50077.

64 Salois MN, Gugger JA, Webb S, Sheldon CE, Parraga SP, Lewitt GM et al. Effects of TP63 mutations on keratinocyte adhesion and migration. Exp Dermatol 2023; 32: 1575–1581.

65 Liu X, Ory V, Chapman S, Yuan H, Albanese C, Kallakury B et al. ROCK inhibitor and feeder cells induce the conditional reprogramming of epithelial cells. Am J Pathol 2012; 180: 599–607.

66 Suprynowicz FA, Upadhyay G, Krawczyk E, Kramer SC, Hebert JD, Liu X et al. Conditionally reprogrammed cells represent a stem-like state of adult epithelial cells. Proc Natl Acad Sci U S A 2012; 109: 20035–20040.

67 Hynds RE, Butler CR, Janes SM, Giangreco A. Expansion of Human Airway Basal Stem Cells and Their Differentiation as 3D Tracheospheres. Methods Mol Biol 2019; 1576: 43–53.

68 Kassis T, Hernandez-Gordillo V, Langer R, Griffith LG. OrgaQuant: Human Intestinal Organoid Localization and Quantification Using Deep Convolutional Neural Networks. Sci Rep 2019; 9: 12479.

69 Patel H, Manning J, Ewels P, Garcia MU, Peltzer A, Hammarén R et al. nf-core/rnaseq: nf-core/rnaseq v3.24.0 - Selenium Seahorse. doi:10.5281/zenodo.19486760.

70 Di Tommaso P, Chatzou M, Floden EW, Barja PP, Palumbo E, Notredame C. Nextflow enables reproducible computational workflows. Nat Biotechnol 2017; 35: 316–319.

71 Soneson C, Love MI, Robinson MD. Differential analyses for RNA-seq: transcript-level estimates improve gene-level inferences. F1000Res 2015; 4: 1521.

72 Love MI, Huber W, Anders S. Moderated estimation of fold change and dispersion for RNA-seq data with DESeq2. Genome Biol 2014; 15: 550.

73 Zhu A, Ibrahim JG, Love MI. Heavy-tailed prior distributions for sequence count data: removing the noise and preserving large differences. Bioinformatics 2019; 35: 2084–2092.

74 org.Hs.eg.db. Bioconductor. http://bioconductor.org/packages/org.Hs.eg.db/ (accessed 13 Apr2026).

75 Hervé Pagès, Marc Carlson, Seth Falcon, Nianhua Li. AnnotationDbi. Bioconductor, 2017 doi:10.18129/B9.BIOC.ANNOTATIONDBI.

76 Durinck S, Spellman PT, Birney E, Huber W. Mapping identifiers for the integration of genomic datasets with the R/Bioconductor package biomaRt. Nat Protoc 2009; 4: 1184–1191.

77 Bharat A, Querrey M, Markov NS, Kim S, Kurihara C, Garza-Castillon R et al. Lung transplantation for patients with severe COVID-19. Sci Transl Med 2020; 12. doi:10.1126/scitranslmed.abe4282.

78 Chua RL, Lukassen S, Trump S, Hennig BP, Wendisch D, Pott F et al. COVID-19 severity correlates with airway epithelium-immune cell interactions identified by single-cell analysis. Nat Biotechnol 2020; 38: 970–979.

79 Loske J, Röhmel J, Lukassen S, Stricker S, Magalhães VG, Liebig J et al. Pre-activated antiviral innate immunity in the upper airways controls early SARS-CoV-2 infection in children. Nat Biotechnol 2022; 40: 319–324.

80 Melms JC, Biermann J, Huang H, Wang Y, Nair A, Tagore S et al. A molecular single-cell lung atlas of lethal COVID-19. Nature 2021; 595: 114–119.

81 Trump S, Lukassen S, Anker MS, Chua RL, Liebig J, Thürmann L et al. Hypertension delays viral clearance and exacerbates airway hyperinflammation in patients with COVID-19. Nat Biotechnol 2021; 39: 705–716.

82 Yoshida M, Worlock KB, Huang N, Lindeboom RGH, Butler CR, Kumasaka N et al. Local and systemic responses to SARS-CoV-2 infection in children and adults. Nature 2022; 602: 321–327.

83 Ziegler CGK, Miao VN, Owings AH, Navia AW, Tang Y, Bromley JD et al. Impaired local intrinsic immunity to SARS-CoV-2 infection in severe COVID-19. Cell 2021; 184: 4713–4733.e22.

84 Luke Zappia AL. zellkonverter. Bioconductor, 2020 doi:10.18129/B9.BIOC.ZELLKONVERTER.

85 Amezquita RA, Lun ATL, Becht E, Carey VJ, Carpp LN, Geistlinger L et al. Orchestrating single-cell analysis with Bioconductor. Nature Methods 2019; 17: 137–145.

86 Hao Y, Stuart T, Kowalski MH, Choudhary S, Hoffman P, Hartman A et al. Dictionary learning for integrative, multimodal and scalable single-cell analysis. Nat Biotechnol 2024; 42: 293–304.

